# Population context drives cell-to-cell variability in interferon response in epithelial cells

**DOI:** 10.1101/2023.05.22.541682

**Authors:** Camila Metz-Zumaran, Patricio Doldan, Francesco Muraca, Yagmur Keser, Pascal Lukas, Benno Kuropka, Leonie Küchenhoff, Soheil Rastgou Talemi, Thomas Höfer, Christian Freund, Elisabetta Ada Cavalcanti-Adam, Frederik Graw, Megan Stanifer, Steeve Boulant

**Affiliations:** Department of Molecular Genetics and Microbiology, University of Florida, College of Medicine, 1200 Newell Drive, 32610 Gainesville, Florida, USA; Department of Infectious Disease, Virology, University Hospital Heidelberg, Im Neuenheimer Feld 344, 60120 Heidelberg, Germany; BioQuant-Center for Quantitative Biology, Heidelberg University, 60120 Heidelberg, Germany; Institute of Chemistry and Biochemistry, Protein Biochemistry, Freie Universität Berlin, Thielallee 63, 14195, Berlin, Germany; Laboratory of Systems Pharmacology, Department of Systems Biology, Harvard Medical School, Boston, Massachusetts, USA; Division of Theoretical Systems Biology, German Cancer Research Center (DKFZ), Heidelberg, Germany; Max Planck Institute for Medical Research, Heidelberg, Germany; Cellular Biomechanics, University of Bayreuth, Bayreuth, Germany; Interdisciplinary Center for Scientific Computing, Heidelberg University, Heidelberg, Germany; Department of Medicine 5, Friedrich-Alexander-Universität Erlangen-Nürnberg, 91054 Erlangen, Germany

## Abstract

Isogenic cells respond in a heterogeneous manner to interferon. Using a micropatterning approach combined with high-content imaging and spatial analyses, we characterized how the population context (position of a cell with respect to the neighboring cells) of human intestinal epithelial cells affects single cell response to interferons. We identified that cells at the edge of a cellular colony are significantly more responsive than cells embedded within this colony. We determined that this spatial heterogeneity in IFN response was the result of the polarized basolateral distribution of the IFN receptors making cells located in the center of a cellular colony not responsive to ectopic IFN stimulation. We could demonstrate that this population context driven cell-to-cell variability influences the outcome of viral infection as cells embedded in a cellular colony are not protected by interferons and therefore more susceptible to infection. Our data highlights that the behavior of individual isolated cells does not directly translate to their behavior in a population, placing the population context as a key driver of cell-to-cell heterogeneity in IFN response.

## Introduction

Interferons (IFNs) are the first line of antiviral innate immune defense. There are three types of IFNs, type I, II, and III. While type II IFNs are mostly produced by immune cells (1, 2), type I and type III IFNs are produced by all cell types. Type I IFNs and type III IFNs bind to the heterodimeric receptors IFN-alpha receptor (IFNAR) IFNAR1/IFNAR2 (3) and IFN-lambda receptor (IFNLR) IFNLR1/IL10Rβ (4, 5), respectively. The IFNLR1 subunit of the type III IFN receptor is mostly expressed in epithelial cells and in some immune cells conferring the type III IFNs a key role to protect mucosal surfaces against viral infection (6–8). In response to virus infection, IFNs are produced and secreted from infected cells and bind to their respective receptors inducing the activation of the Janus kinase (JAK)-Signal Transducer and Activator of Transcription Proteins (STAT) signaling pathway (9). Following activation of STAT1 and STAT2 via phosphorylation, these proteins associate with Interferon Regulatory Factor 9 (IRF9) to form the Interferon Stimulated Gene Factor 3 (ISGF3) complex, which translocates into the nucleus, leading to transcription of interferon stimulated genes (ISGs) that combat viral replication and spread (10, 11).

Most studies aiming at understanding regulation of signal transduction during IFN-mediated signaling have classically used bulk analysis approaches, where the measured parameters represent an average of an entire cell population. For example, when monitoring the kinetics of STAT1/STAT2 activation following IFN treatment, bulk approaches will only provide the time for STAT1/STAT2 to be phosphorylated within the cell population (average phosphorylation time). This does not provide information related to the proportion of cells that responded to IFNs and activated STAT1/STAT2 and similarly, does not address whether all cells responded with the same kinetics to the IFN treatment. Recent studies have discovered cell-to-cell variability to be a central feature of cell populations, even for genetically identical cells growing in the same environment (12). This cell-to-cell variability in response to various stimuli is often referred to as single cell heterogeneity within a cell population. The effects of single cell heterogeneity are wide-ranging, and affect central cellular pathways (13, 14), phenotypic outcomes (15, 16), and even drug sensitivity (17, 18). Intriguingly, with respect to viral infection and IFN-mediated signaling, previous studies have demonstrated a large-scale heterogeneity in the decision and timing of individual cells to produce IFNs upon a pathogenic trigger (19–23). In these studies, virus infection or treatment with synthetic pathogen surrogates only lead to expression of IFNs in a certain percentage of cells, while a subpopulation remained unresponsive, even if proven that the unresponsive cells were infected/stimulated. Additionally, the antiviral state induced by type I IFN was shown to be heterogeneous within one cell population, as IFN treatment did not induce expression of protective ISGs and an antiviral state in a fraction of cells independent of the IFN dosage (20, 21, 23–25). In these studies, which evaluated the heterogeneity during IFN expression and downstream signaling, it has been suggested that the mechanism behind the cell-to-cell variability is attributed to stochastic events along the IFN signal transduction pathways. Stochastic events are a probabilistic distribution of behavior rather than deterministic phenotypes regulated by the molecular machinery. Stochastic events arise from ‘noise’, a term describing some randomness of molecular interactions in the cellular environment (26). This “noise” has been proposed to be critical in cellular decision making and determines, for example, if a cell responds to IFN or not (20, 21, 23, 27).

An alternative explanation for the origins of this cell-to-cell variability is that the molecular machinery instead of stochastic events regulate/drive the heterogeneity of response (i.e. deterministic view) (28). With novel methodologies that allowed elucidating highly complex networks, apparent stochastic events could later be explained by newly discovered mechanisms (29–31). A major determinant for cell-to-cell variability in adherent cell culture systems, that is rooted on tight regulation by the cellular machinery, is the population context (28). The parameters that constitute the population context of an individual cell are the local cell density, cell-to-cell contacts, and relative location within the population. Various molecular mechanisms sense these parameters and translate them to a population-dependent behavior including changes in polarization state, proliferation rate, sensitivity to apoptosis, metabolic state, and cell motility (28). Population-dependent behavior thereby could shape the distribution of single-cell phenotypic properties, leading to the population heterogeneity in genetically identical cells. The population heterogeneity has a large impact on molecular and cell biology. Snijder et al. (32) showed that virus infectivity, endocytic events, and cellular lipid composition were determined by adaptation of single cells to their population context. It was further demonstrated that cell confluence, a central parameter of the population context, induces major changes at the molecular level, leading to differential lipid distribution (33) or protein expression (34) when cells are grown at high vs. low density. Taking together the knowledge obtained from these studies on the population context, it is conceivable that IFN-dependent signaling in isogenic populations is also affected by the population context. We aim to address which role the population context plays on the heterogenous IFN-dependent response in a cell population in adherent intestinal epithelial cells.

The intestinal epithelium separates host tissue from microbiota, and is required to act as a barrier to maintain homeostasis. Intestinal epithelial cells (IECs) polarize and are organized in an impenetrable monolayer. Adjacent cells form junctional complexes as intercellular attachment structures which prevents molecule diffusion (35). This results in a highly dense tissue, in which microbiota is in contact with the apical membrane of IECs and cannot trespass to the *lamina propria*, which is in contact with the basolateral side of IECs. Therefore, *in vivo*, the population context of IECs is characterized by high local density, polarization, and cells being embedded in a monolayer.

Despite the extensive study of antiviral innate immunity in the intestinal epithelium as part of the mucosal barrier, little emphasis has been put on understanding how the population context affects the single cell response to IFNs. Interestingly, it was reported that treatment of a clonal population of mouse derived IECs with type I or type III IFNs induced a heterogeneous response characterized by a responder and a non-responder sub-population independent on the cytokine concentration (36). This heterogeneity in IFN response was also seen in human IECs, where even at very high concentrations type III IFNs were never able to fully protect all cells from virus infection while type I IFN was (37). In our study, we aim to better understand the origins of cell-to-cell variability during an IFN-dependent immune response in human IECs. We combined spatial and quantitative analysis of IFN-mediated signaling with micropatterning approaches to address how the population context impacts response of IECs to IFN treatment. Micropatterning of defined adhesion areas for cell populations allows to tune geometry and size while keeping control over cell density (38). We observed that only a fraction of the cells seeded in partially confluent monolayer responded to apical IFN treatment, and that responsive cells were positioned at the edge of the cell population. Accordingly, cells seeded in a confluent monolayer were less responsive to IFNs as compared to sparsely seeded cells, in which most of the cells had no neighbors and had a similar population context to cells located at edges of a population. This spatial regulation of IFN response disappeared when cells were treated from the basolateral side, which we identified was due to a polarization of the IFN receptor. Furthermore, using IECs which would not form tight barriers, we could demonstrate that the heterogeneity in IFN response during apical treatment is caused by restricted accessibility of the IFN to the respective receptor. Together our results highlight that the population context can greatly impact IFN signaling and is a key parameter to consider when performing experiments in polarized cells.

## Results

### Cell location within a population influences its response to IFN

Previous work has shown that within a cell population, a fraction of the cells do not respond to IFNs despite being genetically identical to the responding cells and having fully functional signal transduction pathways downstream the receptors (19–23). To address whether the population context (i.e., location of a cell within a population) can modulate IFN-mediated signaling, we exploited our previously described human-colon carcinoma T84 cells expressing a fluorescent protein (fp) under the transcriptional control of the interferon stimulated gene (ISG) MX1 promoter (T84-prom-Mx1-fp) (39). With this reporter system, cells are only fluorescent upon IFN-mediated signaling (Supp. Fig. 1), thereby allowing the visualization of the response of each individual cell within a population. T84-prom-Mx1-fp cells seeded at a medium density were mock-treated or treated with type I or III IFNs for 24 h before fixation and analysis using fluorescence microscopy. When seeded at this density, IECs often form small cellular colonies, instead of attaching to the substrate as individual cells uniformly distant from each other (Fig. 1A). This property is likely due to the intrinsic function of IECs to form a cellular epithelial monolayer through tight junction formation. Interestingly, analysis of the location of the cells that became fluorescent upon IFN treatment revealed that mostly isolated cells (cells lacking neighboring cells) and cells located at the edge of a small cellular colony respond to IFNs (Fig. 1A, yellow arrows). In contrast, cells in the center of a colony remained unresponsive (Fig. 1A, red arrows). To quantify how the location of a cell within a population impacts its response to IFN, we performed an unbiased analysis of the cell positioning using DBSCAN-CellX (URL: https://github.com/GrawLab/DBSCAN-CellX/). DBSCAN-CellX is a density-based clustering algorithm allowing us to determine the spatial distribution and positioning of cells in a 2D plane. In brief, our analytic pipeline allows for the registration of the XY-coordinates of each individual cell and for the unbiased determination of whether cells are located at the edge or the center of a colony (Fig. 1B). Additionally, the relative location of an individual cell with regard to the edge or center of a cell population is quantified by the “edge degree”, which represents the distance of a cell from the edge of its colony. The higher the edge degree, the larger the distance from the edge. Cells at the edge are defined by an edge degree of 1, while an edge degree of 0 represents single cells that have no neighbors.

**Figure 1:**
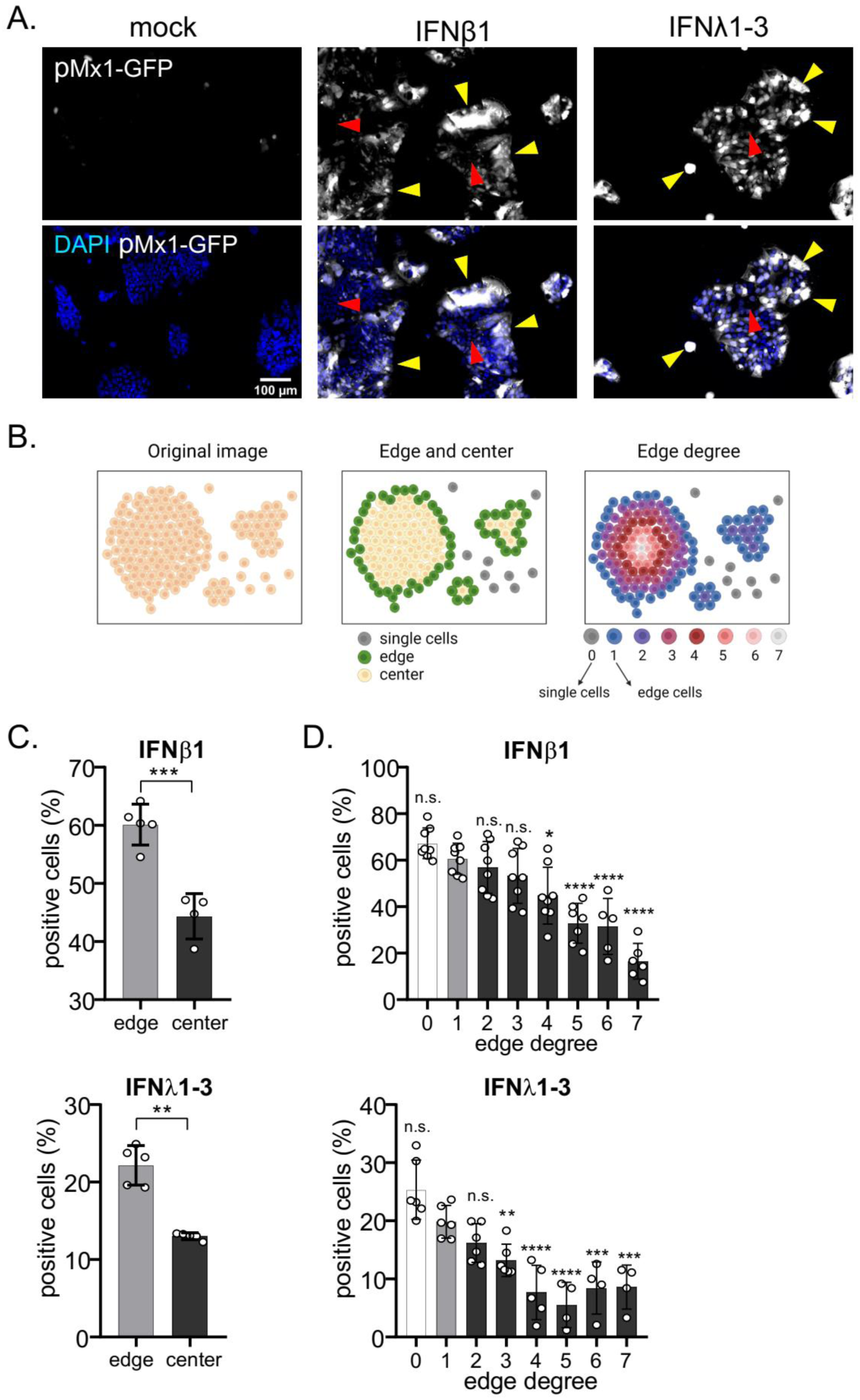
Location of a cell within a population determines its responsiveness to IFN treatment. T84-prom-Mx1-fp seeded at medium density were mock treated or treated with 2000 IU/mL IFNβ1 or 300 ng/mL IFNλ1-3 for 24 h. Cell nuclei were stained with DAPI and fluorescence microscopy was performed. (A) Representative images of cells (nuclei stained with DAPI are blue) expressing the fluorescent reporter (white). Yellow arrows point at IFN responder single cells or cells located at the colony edges. Red arrows point at non-responder cells in the colony center. (B) Correlation between single cell location and IFN-responsiveness was assessed using DBSCAN-CellX. Schematics depicting how the tool annotates cells according to their location at the edge or at the center of a cluster, or according to their edge degree are shown. (C, D) Quantification of the percentage of positive fluorescent cells as compared to mock-treated cells. (C) Edge vs. center cells. (D) Percentage of positive fluorescent cells dependent on the edge degree. An edge degree of 0 define single cells (no neighbors) and edge degree 1 are cells at the border of a colony. The higher the edge degree, the larger the distance from the edge. Error bars indicate standard deviations. n ≥ 3 biological replicates. n.s. =not significant. P<0.05 *, P<0.01 **, P<0.001 ***, P <0.0001 **** as determined by (C) Unpaired t test with Welch’s correction, and (D) ordinary one-way ANOVA with Dunnett’s multiple comparison test using edge degree 1 as reference.

Analysis of the location of the IFN responsive cells within the cell population using the DBSCAN-CellX algorithm revealed that, independent of whether cells were treated with type I or type III IFNs, a significantly higher percentage of edge cells responded to IFNs as compared to center cells (Fig 1C, type I IFN or IFNβ1, top panel; type III IFN or IFNλ1-3, bottom panel). Analysis of the single cell IFN-dependent signaling in correlation to the edge degree showed that the most responsive cells are those lacking neighboring cells (edge degree 0) (Fig. 1D). Importantly, we observed a negative correlation between the edge degree and the percentage of positive cells, further reinforcing that cells located within a population are less responsive to both, type I and type III IFNs (Fig. 1D). Altogether, correlating IFN responsiveness of individual cells to their locations within a population suggests a heterogeneous immune response, in which cells at the edge of cellular colonies are overall more responsive than cells situated inside the colony.

### Micropatterning results in standardized IEC populations, revealing spatial segregation of immune signaling

To fully address whether there is a correlation between a cell location within its population and the extent by which it responds to IFN, we need to employ standardized methods that allow us to control how many cells in a population are located at the center or edge of a cellular colony. For this, we exploited a micropatterning method enabling us to create cell populations of defined and uniform sizes. For this method, a glass surface is passivated with PLL-PEG, an antifouling agent to which cells cannot adhere (Fig. 2A). A Quartz-Mask imprinted with transparent patterns is then overlaid on the passivated surface and illuminated with UV-light in the presence of ozone. As Quartz reflects light, the UV-light can only pass through the transparent areas, thereby depleting the PLL-PEG at discrete locations creating size- and shape-specific patterns on which cells can grow. Using this approach, cells can be grown on controlled micropatterns that provide cells with the same population context.

**Figure 2:**
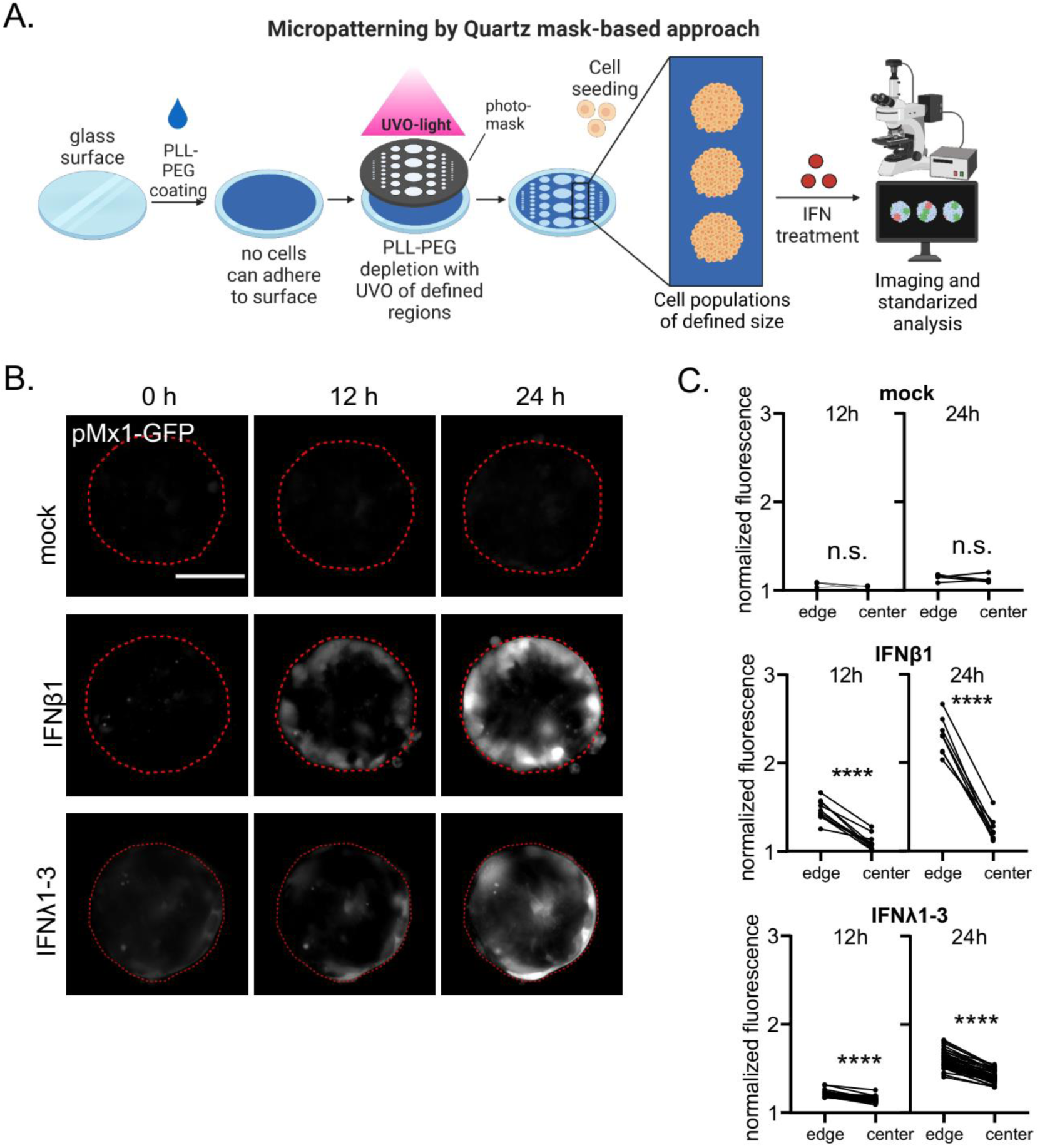
Cells located in the center of a cellular colony are non-responsive to IFNs. (A) Schematic depicting the glass micropatterning approach using a Quartz-mask. (B, C) T84-prom-Mx1-fp cells seeded on circular micropatterns (200 µm diameter) were mock treated or treated with 2000 IU/mL IFNβ1 or 300 ng/mL IFNλ1-3. Fluorescent imaging was performed at 0 h, 12 h, and 24 h post treatment. (B) Representative images. The red line represents the edge of the patterns. Expression of the fluorescent reporter is depicted in white. Scale bar = 100µm. (C) The reporter expression for each single population was quantified by measuring the mean fluorescence intensity (MFI) at the edge and the center of a population at 12 h or 24 h post treatment, and normalizing it to the corresponding population 0 h post treatment. Each dot is one cell population (seeded on one micropattern), lines connect edge and center of the same cell population. n ≥ 3 biological replicates. n.s. =not significant, P <0.0001 **** as determined by Paired t test.

To address the response of IECs to IFNs in these homogenous cell populations, T84-prom-Mx1-fp cells were seeded on micropatterned glass and were treated with type I or III IFNs. Response to IFNs was addressed by fluorescence microscopy at 0, 12, and 24 h post-treatment. Results showed that mostly cells located at the edge of the pattern responded to IFNβ1 treatment (Fig. 2B). This was supported by the quantification of the mean fluorescent intensity (MFI) of the cells located at the edge and center of the population. Edge cells were significantly more fluorescent after type I IFN treatment as compared to center cells. Of note, center cells maintained similar values to mock-treated cells (Fig. 2C) suggesting that in the center of a cell population IECs barely respond to type I IFN treatment. IFNλ1-3 treatment also induced a stronger response at the edge as compared to the center of the micropatterns(Fig. 2B-C). Contrary to the IFNβ1 treatment, center cells were also responsive to IFNλ1-3, however, significantly less than edge cells (Fig. 2B-C). Together, our data strongly suggest that IECs located at the edge of a cell population respond more efficiently to IFNs compared to cells embedded within the cell population.

### Cellular density impacts response of IECs to interferon treatment

To address which part of the IFN-mediated signaling pathway is impaired in cells located at the center of a population, we compared the response of T84 cells seeded at high (205,000 cells/cm^2^) *vs.* low (27,000 cells/cm^2^) cellular density. At high density, cells form a continuous intact monolayer, in which each cell is in contact with neighboring cells from all sides, representing the center of a population (Fig. 3A). On the contrary, at low cell density most of the cells can be considered “edge” cells as they are isolated or are part of small cellular colonies with at least one side lacking a neighboring cell (Fig. 3A). IECs seeded at high and low density were treated with IFNβ1 or IFNλ1-3 (Fig. 3A). The IFN-mediated expression of the ISGs IFIT1, Mx1, and Viperin were measured overtime post-IFN treatment using reverse transcriptase quantitative PCR (RT-q-PCR) (Fig. 3B). Analysis of ISG mRNA expression levels in IFN treated cells normalized to mock treated cells revealed that cells at low density induce significantly higher ISG transcription compared to cells seeded at high density, with almost no transcriptional upregulation of ISG in cells seeded at high densities (Fig. 3B). These results were confirmed using our T84-prom-Mx1-fp reporter cell lines. T84 prom-Mx1-fp cells expressing an H2B-turquoise plasmid (to visualize cell nuclei) were seeded at high and low density and treated 24 h post-seeding with increasing concentrations of IFNβ1 or IFNλ1-3. Following treatment, single cell expression of the fluorescent reporter was followed using live fluorescence imaging for 24 h. Interestingly, we observed a dose dependent response to IFN treatment in cells seeded at low density but not in cells seeded at high density (Supp. Fig. 2). Importantly, cells seeded at high density only responded minimally to IFN treatment (Supp. Fig. 2). These results mirror the intrinsic ISG expression levels as measured using RT-q-PCR (Fig. 3B) further demonstrating that the population context significantly impacts the response of IECs to IFN treatment.

**Figure 3:**
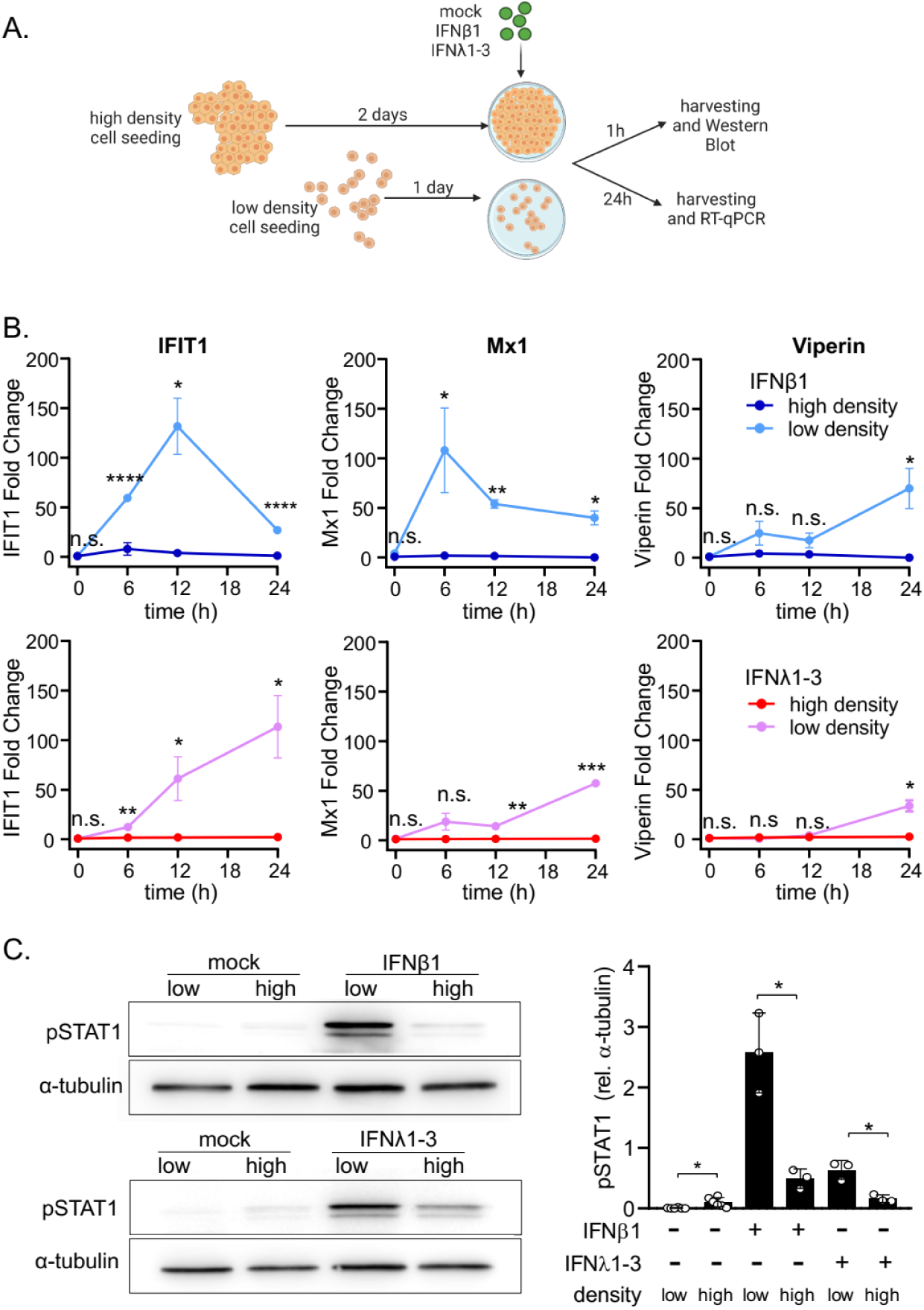
Cell density negatively correlates with IFN-dependent signaling. T84 cells seeded at high and low density were mock treated, or treated with 2000 IU/mL IFNβ1 or 300 ng/mL IFNλ1-3. (A) Schematic depicting the experimental setup. (B) 24 h post IFN treatment, RNA was harvested to evaluate the transcription of the representative ISGs IFIT1, Mx1, and Viperin using q-RT-PCR. ISG relative expression was normalized to the mock-treated cells of the respective time-point (fold change). (C) 1 h post treatment, cellular protein extracts were collected to assess the phospho-STAT1 (pSTAT1) abundance by Western Blot. pSTAT1 was quantified relative to the housekeeping protein α-tubulin. (B, C) n = 3 biological replicates. n.s. =not significant. P<0.05 *, P<0.01 **, P<0.001 ***, P <0.0001 **** as determined by Unpaired t test with Welch’s correction.

The first step in IFN-mediated signaling is binding of IFN to its receptor, which induces the activation of JAK1 that in turn phosphorylates STAT1/STAT2 (10). To address whether cell density can impact the phosphorylation of STATs following IFN stimulation, T84 cells seeded at low and high densities were stimulated with either type I IFN (IFNβ1) or type III IFN (IFNλ1-3). 1 h post-stimulation, the phosphorylation status of STAT1 was addressed by Western Blot analysis. Results show that treatment of IECs seeded at low density with either type of IFN induced significantly higher STAT1 phosphorylation as compared to cells seeded at high density (Fig. 3C).

Altogether, we observe a negative correlation between IEC density and response to IFNs: confluent cells are almost unresponsive to IFNs while sparse cells show high levels of STAT1 phosphorylation and downstream ISG expression upon IFN treatment.

### Basolateral treatment of IECs with IFN suppresses the spatial heterogeneity of IFN-mediated signaling

IECs are polarized, meaning that they have both an apical and a basolateral membrane (Fig. 4A). The basolateral side represents the bottom of the cell, which is in contact with the cell culture vessel *in-vitro* and in contact with the lamina propria *in-vivo*. The apical membrane represents the top of the cell facing the cell culture medium *in-vitro* and the lumen of the gut *in-vivo*. A possible explanation to account for the greater response to IFNs of IECs located at the edge of a population or of IECs seeded at low density could be if the IFN receptors are mostly localized on the basolateral side of IECs. IFN treatment of cells seeded on glass or plastic surfaces would not lead to IFN-mediated signaling for the cell located at the center of a population or in a confluent monolayer as IFNs would not be able to access the basolateral side of the cells where the IFN receptors may be localized. On the contrary, IFNs can stimulate cells located at the edge of a colony as edge cells are not polarized and do not show an asymmetric basolateral distribution of their receptors (40). To directly challenge this model, we seeded T84 cells on transwell inserts to allow for the formation of a polarized cell monolayer characterized by formation of tight junctions. Tight junctions are intercellular adhesion complexes in epithelia that control paracellular permeability. They form an adhesive belt between epithelial cells that control the diffusion of molecules between cells. Three major transmembrane proteins (occludin, claudins and junction adhesions molecule (JAM)) interact with the neighboring cells, and associate with the intracellular protein Zonula Occludens 1 (ZO1). ZO1 therefore has a central role within junctional complexes, acting as a scaffold protein that anchors the actin cytoskeleton with the tight junction transmembrane proteins (41). Polarization and formation of tight junctions preventing diffusion of molecules across the epithelium barrier was controlled by immunofluorescence staining of ZO1, monitoring of the trans-epithelial electrical resistance (TEER), and restriction of FITC-dextran free diffusion from the apical to the basolateral transwell compartment (Supp. Fig. 3). As expected, T84 cells grown on transwell inserts formed a tight junction belt between individual epithelial cells (Supp. Fig. 3A), established a TEER-value characteristic of polarized T84 cells (Supp. Fig. 3B) (42), and formed a tight monolayer of cells that prevents diffusion of molecules between cells (Supp. Fig. 3C). Polarized IECs on transwell inserts were treated with IFNs either from the apical or basolateral side for 24 h (Fig. 4A, left panel). Quantitative-RT-PCR analysis of the expression of the ISG IFIT1 showed that basolateral treatment induced a significantly higher response as compared to apical treatment (Fig. 4A, right panel). Together these findings suggest that the IFN receptors might be enriched at the basolateral side of polarized T84 cells.

**Figure 4:**
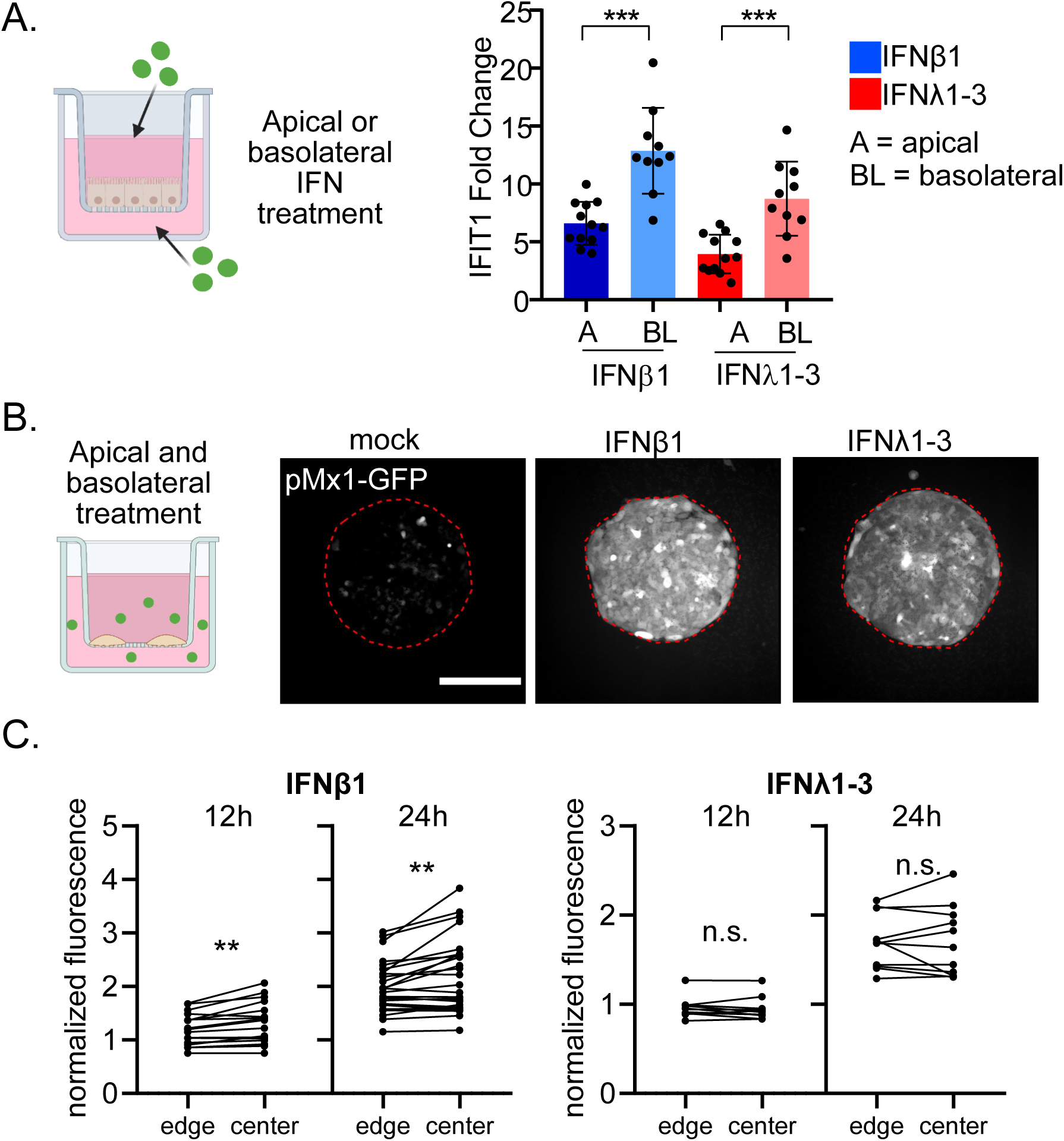
T84 cells better respond to IFN when stimulated on their basolateral side. (A) T84 cells seeded on transwell inserts were mock treated, or treated from the apical (A) or basolateral (BL) side with 2000 IU/mL IFNβ1 or 300 ng/mL IFNλ1-3. 24 h post treatment, RNA was harvested, and q-RT-PCR was used to evaluate the expression of the ISG IFIT1. Data is normalized to mock (fold change). (B, C) T84 prom-Mx1-fp cells were seeded on micropatterned transwell membranes. Cells were mock treated, or treated simultaneously from the apical and basolateral side with 2000 IU/mL IFNβ1 or 300 ng/mL IFNλ1-3. Cells were fixed at 0 h, 12 h, and 24 h post treatment and fluorescent imaging was performed. (B) Representatives images showing treated T84 cell populations. The red line represents the edge of the patterns. Expression of the fluorescent reporter is depicted in white. Scale bar=100µm. (C) The reporter expression was quantified by measuring the mean fluorescence intensity (MFI) at the edge and the center of a population at 12 h or 24 h post treatment, and normalizing it to the corresponding 0 h post treatment at the edge and center, respectively. Each dot is one cell population (seeded on one micropattern), lines connect edge and center of the same cell population. (A, C) n ≥ 3 biological replicates. n.s. =not significant, P<0.01 **, P<0.001 *** as determined by (A) Unpaired t test with Welch’s correction, and (C) Paired t test.

To address whether basolateral treatment of cells seeded on micropatternscan render the center cells responsive to IFNs, we micropatterned transwell inserts (Supp. Fig. 4) and seeded our T84-prom-Mx1-fp reporter cell line on them. In this setup, T84-prom-mx1-fp cells are simultaneously treated with IFNs from the apical and basolateral side (Fig. 4B). Analysis using fluorescent microscopy revealed that the prom-Mx1-fp reporter expression was not restricted to the edge cells anymore, but was instead also found at the center of the patterns (Fig. 4B). Quantification of the fluorescent signal relative to mock treated cells showed that center cells are inducing higher ISG levels compared to edge cells following type I IFN treatment (Fig. 4C), which is opposite to when micropatterned cells are stimulated with IFNs only from their apical side (Fig. 2B-C). Altogether, our data show that the spatial restriction of immune response following IFN treatment can be bypassed by stimulating cells from their basolateral side.

### IFNAR2 and IL10RB are predominantly localized at the basolateral side of polarized T84 cells

To directly address whether the IFN receptors are asymmetrically distributed in polarized T84 cells and located at their basolateral side, T84 cells were grown as a monolayer on transwell inserts. Apical or basolateral surface proteins were biotinylated by addition of cell non-permeable reactive NHS-biotin to the apical or basolateral compartment of the transwell inserts, respectively. Biotinylated proteins were pulled down using streptavidin beads and identified using mass spectrometry (Fig. 5A). Mass spectrometry results showed a significant enrichment of biotinylated surface proteins, while non-specific binding to beads was minimal (Fig. 5B), thereby confirming efficiency of the pulldown. We controlled the specificity of our assay by determining the correct localization of known apical (ALPP and ALPPL2) and basolateral (ATP1B1 and ATP1A1) polarized intestinal epithelial cells markers (Fig. 5C, highlighted in green). Analysis of the surface proteome of the apical and basolateral membranes of IECs revealed that both, IFNAR2 and IL10RB, were readily detectable at the surface of T84 cells with a specific distribution of 75% basolateral vs. 25% apical (Fig. 5C). IFNAR1 and IFNLR1 were not detectable using mass spectrometry likely due to their low expression levels. This distribution of IFNAR2 and IL10RB is in agreement with our findings where we stimulated T84s with IFNs from either the apical or basolateral membrane (Fig. 4A). Together our data strongly suggest a model where the spatial restriction of IFN response to cells located at the edge of a cellular colony is due to the distribution of the IFN receptor mostly at the basolateral side of T84 cells.

**Figure 5:**
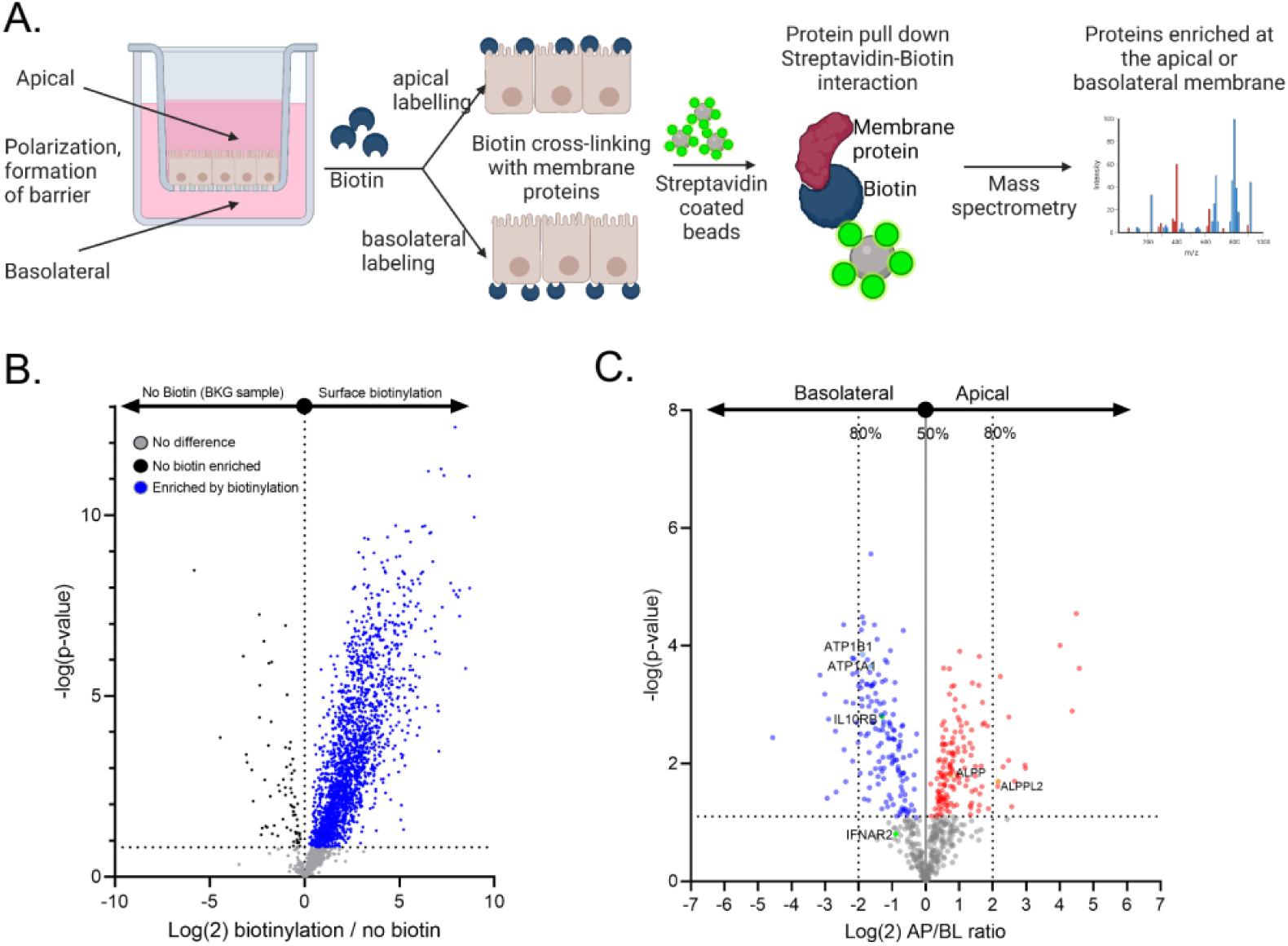
Mass spectrometry of the apical and basolateral proteome confirms the polarized localization of IFN receptors. T84 WT cells were grown as a polarized monolayer on transwell inserts. Apical or basolateral surface proteins were biotinylated by addition of cell non-permeable reactive NHS-biotin to the apical or basolateral compartment of the transwell insert, respectively. Biotinylated proteins were pulled down using streptavidin beads and identified using mass spectrometry. (A) Schematic showing the method. (B) Volcano plot showing enrichment via biotinylation of the surface proteome. (C) Volcano plot showing apical/basolateral log_2_ ratios for detected surface proteins in T84 cells (right). IFNAR2 and IL10BR are both present on the basolateral side of polarized T84 cells cells (n = 6, p-value calculated using FDR <0.01). Known apical and basolateral markers of polarized gut epithelial cells are also highlighted. (B,C) n = 6 biological replicates.

### Tight junctions in polarized T84 cells restrict IFN access to their basolateral receptors

Tight junctions control paracellular permeability, and ZO1 has a central role as a scaffold protein within junctional complexes (41). To address whether tight junctions can prevent the diffusion of IFN to the basolateral side of T84 cells when grown as a polarized monolayer, we created a cell line depleted of the master tight junction protein ZO1. We reasoned that knock-out of ZO1 should disrupt tight junctions, leading to uncontrolled paracellular diffusion between cells, allowing IFN to access the basolateral receptors (Fig. 6A, right panel). Knock-out of ZO1 was validated at the protein level by Western blot analysis (Fig. 6B) and immunofluorescent staining (Fig. 6C). Importantly deletion of ZO1 impaired the formation of a tight monolayer as ZO1 KO cells were significantly delayed in their establishment of a TEER when seeded on transwell inserts as compared to WT cells, and after 5 days post seeding T84 ZO1 KO cells only reached a TEER of 1500 Ω/cm^2^ while T84 WT cells reached a TEER of 6500 Ω/cm^2^ (Fig. 6D). T84 WT and T84 ZO1 KO cells seeded at high (H) and low density (L) were treated apically with IFNs (Fig. 6D). In line with previous results, IFN treatment of sparse T84 WT cells induced significantly higher ISG expression as compared to IFN treatment of confluent T84 WT cells (Fig. 6E). In stark contrast, no difference could be observed on the response to IFNs at low vs. high density in T84 cells depleted of ZO1 (Fig. 6E). These results demonstrate that the tight junction protein ZO1 restricts the paracellular diffusion of IFN between polarized T84 cells, and that access of IFN to the basolateral membrane of a polarized T84 cellular monolayer is a crucial determinant to induce an IFN-mediated response.

**Figure 6:**
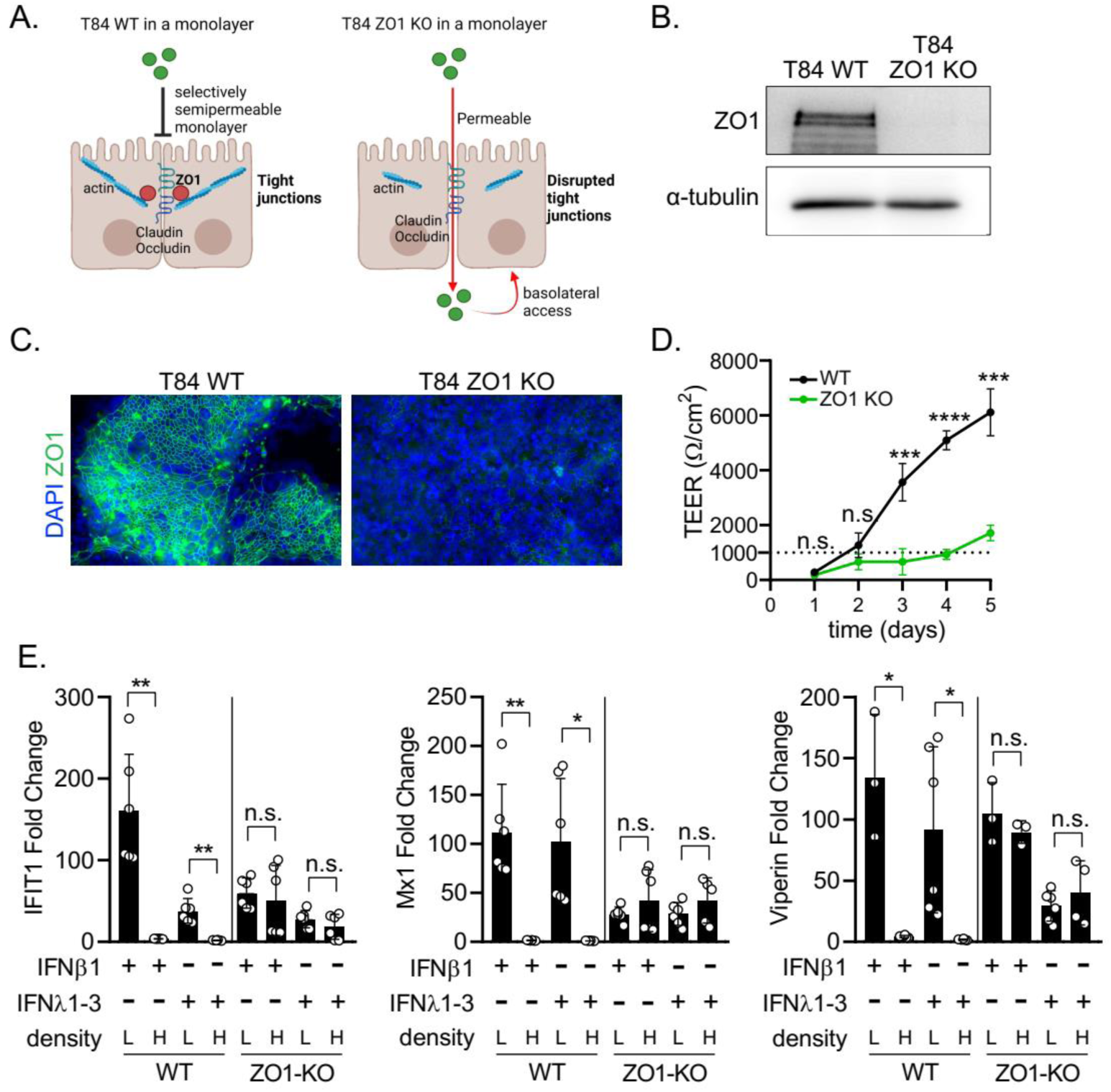
Disruption of tight junctions allows for IFN response in cells at high density. (A) Schematic depicting paracellular diffusion in a monolayer of T84 WT and T84 ZO1 KO cells with disrupted junctional complexes. (B) T84 WT and T84 ZO1 KO cell protein extracts were harvested to control the absence of ZO1 protein in the KO cells by Western Blot. α-tubulin served as a housekeeping protein. Representative image is shown. (C) T84 WT and T84 ZO1 KO cells were fixed and indirect immunofluorescence was performed against the junctional complex protein ZO1 (green). Nuclei were stained with DAPI (blue). Representative image is shown. (D) T84 WT and T84 ZO1 KO cells were seeded on transwell inserts and grown as a polarized monolayer. Transepithelial electrical resistance (TEER) was measured over a period of 5 days. Dotted line shows a TEER of 1000 Ω/cm^2^ corresponding to the resistance formed by confluent polarized T84 cells (48). (E) T84 WT and T84 ZO1 KO cells at high (H) and low (L) density were treated apically with 2000 IU/mL IFNβ1 or 300 ng/mL IFNλ1-3. 24 h post treatment, RNA was harvested, and q-RT-PCR was used to evaluate the expression of the ISG IFIT1. Data is normalized to mock (fold change). (D, E) n ≥ 3 biological replicates. n.s. = not significant, P<0. 05 *, P<0.01 **, P<0.001 ***, P <0.0001 **** as determined by (D) ordinary one-way ANOVA with Dunnett’s multiple comparison test using the positive control as reference, and (E) Unpaired t test with Welch’s correction.

### IEC density significantly affects the IFN-induced protection from virus infection

When performing traditional 2D cell culture experiments, seeding densities are chosen traditionally around 70% confluence or slightly adapted to accommodate for extended culturing times. On the contrary, when working with epithelial cells, high cell density is often employed to better mimic the physiological growing conditions of these cells and to induce cell polarization. Given the localization of the IFN receptors at the basolateral side of T84, we wondered whether cellular density can impact the outcome of viral infection during prophylactic treatment of epithelial cells with IFNs.

T84 cells were seeded at high and low density, and pre-treated with IFNβ1 and IFNλ1-3 for 24 h (Fig. 7A). Cells were then infected with two unrelated viruses, Vaccinia virus (VV) or Mammalian Reovirus (MRV), for 16 h and infection levels were assessed by immunofluorescence microscopy (Fig. 7A). Interestingly, IECs seeded at low cell density pre-treated with IFNs were able to control VV infection better than cells treated at high density (Fig. 7B). When quantifying the number of VV-infected cells, we observed that for both, high and low cell density, infection levels were around 50% (Fig. 7C). However, pre-treatment of cells seeded at low density with IFNβ1 strongly reduced the number of VV-infected cells to ~5%, and IFNλ1-3 reduced it to ~20% (Fig. 7C). In contrast, for cells seeded at high density, IFN pre-treatment had no significant effect on VV infection levels when compared to mock-treated infected cells (Fig. 7C). Similar results were observed for MRV infection, in which IFN pre-treatment of cells at low density significantly reduced the number of infected cells as compared to non-treated cells while no protective effect of IFN pretreatment was observed for cells seeded at high density (Fig. 7D). Together, we could show that the accessibility of the IFN receptor affects the antiviral priming of IECs. This has detrimental consequences for experimental outcomes, since cells at low confluence, where an antiviral state was induced, were able to restrict virus infection. On the contrary, cells at high confluence with less receptors accessible on the apical side, induced a lower response to IFN pre-treatment and therefore were not protected from virus infection.

**Figure 7:**
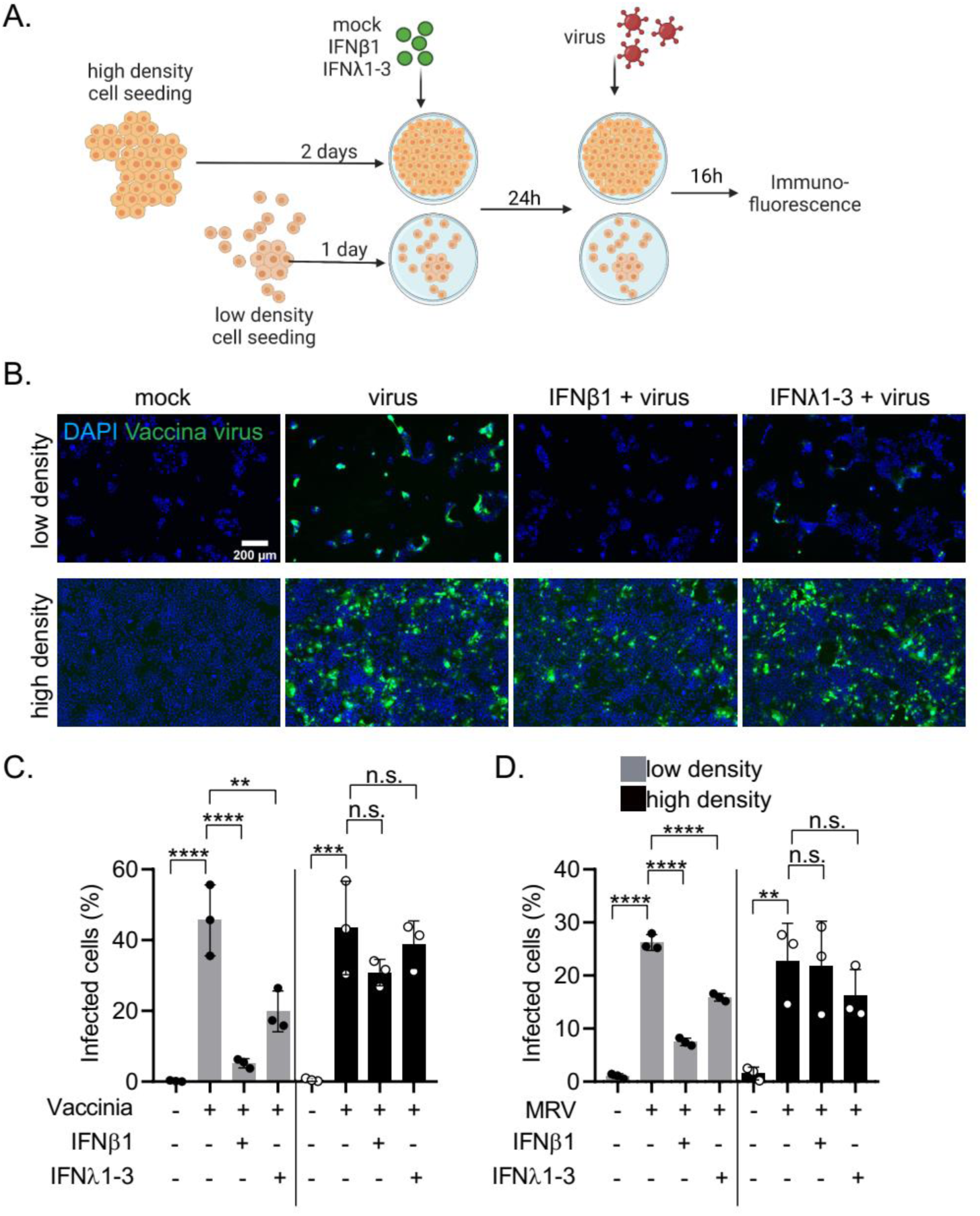
Polarized IFN receptor localization affects induction of an antiviral state in confluent cells. T84 cells seeded at high and low density were mock-treated or pre-treated with 2000 IU/mL IFNβ1 or 300 ng/mL IFNλ1-3. 24 h post treatment, cells were infected with Vaccinia virus-eGFP (VV) or Mammalian Reovirus (MRV) at an MOI of 1 (as determined in T84 WT cells). Infection media was supplemented with the respective IFN. 16 h post infection, cells were fixed, immunostained for viral protein and fluorescence imaging analysis was performed. (A) Schematic of the experimental setup. (B) Representative images showing Vaccinia virus eGFP (green) infected cells. Nuclei were stained with DAPI (blue). Scale bar=200µm. (C, D) Quantification of the number of (C) Vaccinia virus eGFP infected cells and (D) MRV infected cells. n = 3 biological replicates. n.s. = not significant, P<0.01 **, P<0.001 ***, P <0.0001 **** as determined by ordinary one-way ANOVA with Dunnett’s multiple comparison test. Test was performed within high or low density groups, using only virus infected cells (no pretreatment) as reference.

Altogether, our results show that IECs differently respond to IFNs according to their population context. Isolated cells and cells located at the edge of a cellular colony are much more responsive to IFNs compared to cells embedded within a cellular colony. We could show that this spatial restriction of IFN-mediated signaling was due to the basolateral location of the IFN receptors in epithelial cells. Within a cellular colony, central embedded cells only have their apical plasma membrane accessible and as such respond very poorly to IFN treatment. Finally, we could show that this differential response of IECs to IFNs depending on the population context is critical to define whether IECs would be protected or not upon IFN treatment against viral infection. Our work highlights the importance of considering the population context when studying susceptibility of cells to viral infection and efficacy of antiviral measures, as the location of a cell within a population or whether the experimental set up use partially vs. fully confluent cells can severely impact the experimental outcomes.

## Discussion

Determining at the molecular level how IFNs induce a protective antiviral state in IECs is a prerequisite to better understand infectious disease in the gut and to develop novel antiviral therapeutic strategies. Employing single cell and population analysis pipelines, we demonstrated that both type I and type III IFNs induce a heterogeneous response in isogenic human IECs. This response is characterized by cells at the edge of a cellular colony mounting a significantly higher immune response as compared to cells localized in the center of the cell population. We identified that the origin of this cell-to-cell variability is an asymmetric distribution of the IFN-receptors toward the basolateral side of IECs. Cells localized in the center of a colony form a polarized monolayer, and IFNs coming from the cell culture medium (apical side) cannot access the basolateral receptors. On the contrary, cells at the edge of a colony are not polarized and the receptors are localized around the entire cell allowing interaction with IFNs present in the cell culture medium. In accordance with this observation, basolateral IFN treatment induces a homogenous signaling in which all cells, independently of their location within a population (center or edge of colony) respond to IFNs. Importantly, we demonstrated that this polarized IFN-receptor localization can greatly affect the outcome of infection when addressing the protective state induced by IFNs during virus infection. Pre-treatment of confluent IECs with IFNs provide limited protection against viral infection. This finding highlights the importance of considering the population context when studying host cell pathogen interactions and when addressing the potency of the antiviral function of IFNs in epithelial cells.

### Heterogeneous response of cells within a population to IFN treatment

Cell-to-cell variability during IFN-mediated immune responses in isogenic clonal cell populations has been widely observed. However, our understanding of the molecular basis for this cell-to-cell heterogeneity is only at its infancy. It has been reported that individual cells within a population induce ISG expression at different times post type I IFN treatment (23, 24). Moreover, low concentrations of type I IFNs are known to induce a heterogeneous pattern of ISG expression levels with highly responsive, less responsive, and non-responder cell subpopulations (20, 23, 24, 36). Importantly, a similar subpopulation of non-responsive cells was observed in conditions where cells were treated with saturating type I IFN concentrations (36, 43). Altogether this demonstrates that, within a clonal cell population, some cells, although fully equipped at the molecular level to respond to IFNs, are not responsive to these cytokines. This non-responsiveness of a subpopulation of cells to IFNs is not terminally determined. When the non-responder population is isolated and re-treated with type I IFNs, the same heterogeneous pattern of ISG expression (responder and non-responder cells) was induced, thereby excluding the existence of a stable fraction of unresponsive clones (36, 43). This suggests that within a population, cells actively engage in intercellular communication to reprogram cell subpopulations and generate a precise equilibrium between responsive and non-responsive cells.

Using *in-silico* modeling and single cell data, the cell-to-cell variability was explained by stochastic events rooted in ‘biochemical noise’ (20, 24, 36, 43), rather than deterministic events tightly regulated by the molecular machinery. To point out, in these studies single cell behavior was assessed mostly by flow cytometry, in which it is not possible to trace the spatial context of each cell, or fluorescence microscopy without specifically accounting for single cell location (20, 21, 23). In line with previous studies, we also observed that an isogenic population of adherent hIECs treated with either type I or type III IFNs from the apical side results in a strongly heterogeneous immune response, ranging from highly responding to non-responder subpopulations (Fig. 1–2). However, by using tools that integrate the effect of spatial components in cell behavior, we identified that the population context of IECs (position of a cell with respect to its neighbors within a population) explains the heterogeneous response of IECs to IFNs. Instead of a stochastic event driving the heterogeneous response to IFN in IECs, we propose that the polarized distribution of the IFN receptors to the basolateral side of the cells constitutes a deterministic explanation for the observed spatial restriction of IFN response. While we show that basolateral IFN treatment of a polarized monolayer allows for all cells to respond in a population, we still observe a heterogeneous pattern in the amplitude of single cell response (intensity of fluorescent ISG reporter expression (Fig. 4B)), and it remains to be determined to what extent this cell-to-cell variability is stochastic or deterministic. Interestingly, a study by Bhushal et al. (36) also observed the presence of a non-responsive subpopulation upon type III IFN treatment in murine IECs. In this study the number of reactive cells increased upon cell confluence. They further demonstrated that cell polarization and the epigenetic status partially determine the size of the non-responder population, thereby explaining that the heterogeneous response to type III IFNs in mouse IECs is also tightly regulated by the molecular machinery. This and our study thereby highlight the role of cell confluence and the population context during IFN sensing and signaling, which can be incorporated as a deterministic factor and should be re-addressed in previous studies arguing towards a stochastic origin of cell-to-cell variability during IFN signaling.

### Spatial and temporal determinants of cell-to-cell response heterogeneity to extracellular stimuli

Studying cell-to-cell heterogeneity in isogenic populations has been facilitated by single-cell transcriptomic and flow-cytometry, however these methods do not integrate both the spatial and temporal determinants that may characterize responder and non-responder cells. In contrast, high content imaging enables the collection of spatially and temporally resolved data with single cell resolution. Here, we combined high-content imaging using a fluorescent reporter cell line with (a) a bioinformatics method (DBSCAN-CellX) (Fig 1) and (b) a micropatterning method (Fig 2 and 4) to address how the population context (spatial heterogeneity) impacts IFN-mediated immune response and antiviral response in IECs. Using our recently developed DBSCAN-CellX approach (https://github.com/GrawLab/DBSCAN-CellX/), we could quantify the relative location of individual cells within a population with respect to their neighboring cells (Fig 1), providing us with a tool to address single cell behavior in their population context. This allowed us to identify a spatially-dependent heterogeneous response pattern, in which significantly more cells at the edge of a population induced ISG expression as compared to cells in the center of a population. These results were confirmed using our micropatterning approaches that allowed us to create cell populations in which all population context parameters (population size, local density, polarization status) were manipulatable and reproducible (Fig 2 and 4), enabling us to study cell population behavior in an unbiased and controlled manner. Using these tools in combination with high content imaging pipelines promise to improve our understanding of single cell heterogeneity by allowing us to better deconvolve stochastic or deterministic origin of cell-to-cell variability.

### The population context impacts cell ability to mount an antiviral response upon IFN treatment

We demonstrated that cell confluence can greatly affect experimental outcomes while testing the sensitivity of several viruses to IFN treatment. To address whether IFNs are protective against those pathogens, we apically pre-treated cells at high and low density with IFNs prior to virus infection (Fig 7). The conclusions that are drawn from these two different experimental setups (high vs. low cell density) are opposing: Results obtained from low density suggest that IFNs induce a strong antiviral state against the tested viruses. On the contrary, results from confluent cells show that IECs cannot be protected from viruses by IFNs. These experiments highlight the importance of considering cell density when determining the experimental setups, especially in the context of the intestinal epithelium and antiviral immune response. Previous studies demonstrated that cell density is involved in major cellular molecular pathways, and thereby affects lipid composition (33), endocytic events (32) and the expression of central molecules including autophagy markers p62 and LC3II, lysosomal cathepsin D as well as nuclear proteins HDAC1 and Lamin B1 (34). Interestingly, K. Trajkovic et al. (34) treated cells with widely used compounds that lead to undesired changes in cell density, and compared the effect of the compounds to non-density matched or density matched controls, which lead to ambiguous conclusions. This demonstrated that cell density is a potent experimental variable, and they emphasize that a rational experimental design including cell density controls will minimize erroneous interpretation of cell culture data. Importantly, cell density is assumed to be associated with drug resistance and various studies showed that cells embedded in a confluent monolayer are significantly less susceptible to drug treatment (44, 45), which has far reaching effects in the area of drug screening and development within the biomedical industry. We propose that cell density is underestimated during the evaluation of results and must be more actively addressed when planning experiments. Moreover, joined effort must be invested in recognizing population factors involved in biological processes.

### Polarized distribution of IFN receptors

We demonstrate that both type I and type III IFN receptors are enriched on the basolateral membrane of polarized IECs. Polarized IFN-alpha receptor (46) and IFN-gamma receptor (47) localization to the basolateral membrane has been reported before for airway epithelial cells. However, to the best of our knowledge, no report has focused on IFN receptor localization in the gut. A polarized receptor localization might have a physiological relevance, since *in-vivo* IECs are in contact with the *lamina propria* from the basolateral side where immune cells are also situated. On the contrary, the apical membrane faces the gut lumen containing the commensal microbiota system. Sensing IFNs from the sterile basolateral side could be a mechanism to selectively sense IFNs provided by immune cells. IECs also express and secrete IFNs to act in an autocrine and paracrine manner, and to propagate an antiviral immune response. Interestingly and in line with our results, it was demonstrated that after virus infection of polarized hIECs *in-vitro*, IFNλ was secreted predominantly to the basolateral side (48). Further studies must address whether IFN secretion *in-vivo* by IECs occurs on the apical or basolateral side, and how this is relevant in the context of a basolateral IFN receptor localization.

With our study we provide a novel approach to understand the origins of heterogeneity in isogenic populations. We demonstrated that the spatial heterogeneity during IFN response in IECs is originated by a basolateral receptor localization in polarized cells. As our results show, the population context determining the polarized receptor localization can have wide-ranging effects on the experimental outcome, and we suggest that experiments need to be planned accordingly to obtain accurate results.

## Material and Methods

### Cell lines, cell culture media and viruses

HEK293T cells (ATCC) were cultured in Iscove’s Modified Dublecco’s Medium (IMDM) (Gibco #124400-053) supplemented with 10% fetal bovine serum (Sigma Aldrich #12306C) and 1% penicillin/streptomycin (Gibco #15140122). Wild type (WT) T84 (ATCC CCL-248) as well as T84 reporter and knock-out (KO) cells were cultured in a 50:50 mixture of Dulbecco’s modified Eagle’s medium (DMEM) and F12 (Gibco #11320033) supplemented with 10% fetal bovine serum (Sigma Aldrich #12306C) and 1% penicillin/streptomycin (Gibco #15140122).

The IFN-sensing reporter T84 cell lines expressing prom-Mx1-mCherry or prom-Mx1-eGFP were previously generated in our laboratory and described in Doldan et al. (39). The T84 ZO-1 KO cell line was generated using a lentivirus-based CRISPR-Cas9 gene editing system. First, the guideRNA with the sequence gttttagagctagaaatagcaagttaaaataaggctagtccgttatcaacttgaaaaagtggcaccgagtcggtgc was inserted into the plasmid lentiCRISPRv2 containing a Blasticidin resistance. A lentivirus vector system was used to efficiently deliver the CRISPR-Cas9 plasmid to the T84 cells. To first package the plasmid in lentivirus, HEK293T cells at 80% confluence in a 10 cm^2^ dish were transfected with 8 µg of the CRISPR-Cas9 plasmid containing the guideRNA targeting ZO1, 4µg pMDG.2 plasmid and 4µg psPAX plasmid by using the transfection reagent Polyethylenimine (PEI) (Polysciences #23966-100) at a PEI:DNA ratio of 4:1. 3 days post transfection, the supernatant containing lentivirus was collected, spun down to separate it from cell debris at 4000 rcf for 10 mins, and filtered through a 0.45 µm syringe filter (Lab Unlimited #W10462100). To pellet the lentivirus, the supernatant was spun down at 27,000 rpm for 1:40 hours using a SW40 Ti rotor. The lentivirus pellet was resuspended in 100 µL OptiMem (Gibco #31985062) (per yield of one 10 cm^2^ dish) and used for transduction. For transduction, 300,000 WT T84 cells per well in a 6-well plate were treated with 20 µL lentivirus using 3 µL Polybrene transfection reagent (Sigma Aldrich #TR-1003-G) diluted in 3 mL media. After 3 days of incubation, transduced cells were selected with Blasticidin (0.1 mg/mL) (Invivogen #ant-bl-1). Single cell cloning was performed using a limited serial dilution approach to obtain a monoclonal population knocked out for ZO1.

Mammalian Reovirus (MRV) type 3 clone 9 was derived from stocks originally obtained from Bernard N. Fields and was grown and purified by standard protocols (48). Vaccinia virus eGFP is a Western Reserve Vaccinia Virus strain that expresses EGFP under the control of a synthetic Early/Late virus promoter and was first described by J. Mercer and A. Helenius (49). Vaccinia virus eGFP was kindly provided by Jason Mercer and was grown and purified by standard protocols (50).

### Cell culture

Cell seeding on multiwell plates: T84 cells were seeded on rat-collagen (Sigma-Aldrich #C7667-25MG) coated multiwell plates (Corning). For high density, 225,000 T84 cells per well were seeded in 48-well plates. One day post-seeding, medium was exchanged with 0.5 mL fresh culturing medium, and two days post-seeding cells were treated with IFNs. For low density, 30,000 T84 cells per well were seeded in 48-well plates. One day post-seeding cells were treated with IFNs.

Cell seeding on glass bottom 8-well chamber slides (iBIDI): 100,000 T84 cells per well were seeded on glass bottom 8-well chamber slides coated with 2.5% human collagen (Sigma #C5533-5MG) diluted in water. One day post seeding cells were treated with IFNs.

Cell seeding on transwell inserts: 120,000 T84 cells were seeded on rat-collagen (Sigma-Aldrich #C7667-25MG) coated 6.5 mm transwell 3.0 µm Pore Polycarbonate Membrane Inserts (Corning, #3415). Media was exchanged every second day until a polarized cell monolayer was formed. Monolayer permeability and integrity was assessed by measurement of the Transepithelial electrical resistance (TEER) using the EVOM3 Epithelial Volt/Ohm Meter with STX2-PLUS (Word Precision Instruments). When a TEER of ≥ 1000 Ω/cm^2^ was reached, cells were considered polarized forming a tight monolayer.

### Interferon treatment

Human recombinant IFN-beta 1a (IFNβ1) was obtained from Biomol (#86421) and cells were treated with 2000 IU/mL or as described in the figure legend. Human recombinant IFNλ1 (IL-29) (#300-02L), IFNλ2 (IL28A) (#300-2K) and IFNλ3 (IL-28B) (#300-2K) were purchased from Peprotech, and cells were treated by a cocktail of all three type III IFNs in a ratio of 1:1:1, resulting in a final concentration of 300 ng/mL or as described in the figure legend. Cells were treated with IFNs diluted in culturing media (250µL for 48-well plate, 200 µL for Labtec, 200 µL for apical transwell treatment, 800 µL for basolateral transwell treatment, 1 mL for patterned coverslips) and the duration of the treatment is stated in the figure legends.

### Western blot

Cells were harvested and lysed with 1X RIPA buffer (150 mM sodium chloride, 1.0% Triton X-100, 0.5% sodium deoxycholate, 0.1% sodium dodecyl sulphate (SDS), 50 mM Tris, pH 8.0) with cOmplete™ Mini EDTA-free Protease Inhibitor Cocktail (Sigma Aldrich #11836170001) and phosphatase inhibitor PhosSTOP (Millipore Sigma #PHOSS-RO) for 5 min at 37°C. Lysates were collected and protein concentration was measured using the Pierce BCA Protein Assay Kit assay (Thermo Scientific #23225) according to the manufacturer’s protocol. 8 µg protein per condition were separated by SDS-PAGE and blotted onto a 0.2 µm nitrocellulose membrane (Bio-Rad, #1704158) using a Trans-Blot® Turbo™ Transfer System (Bio-Rad). Membranes were blocked with Tris Buffer saline (TBS)-tween (0.5% Tween in TBS) containing 5% Bovine Serum Albumin (BSA) (blocking buffer) for 2 h at room temperature (RT). Primary antibodies against alpha-Tubulin (Sigma #T9026), phospho-STAT1 (BD Transductions #612233) and ZO-1 (Invitrogen #33-9100) were diluted 1:1000 in the same blocking buffer and nitrocellulose membranes were incubated with the antibodies diluted in the blocking buffer overnight at 4°C. Membranes were then washed three times with TBS-T for 5 min at room temperature (RT) while rocking. Anti-mouse antibodies coupled with horseradish peroxidase (HRP) (GE Healthcare #NA934V) were used at 1:5000 dilution in blocking buffer and incubated at RT for 1 h while rocking. Membranes were washed three times with TBS-T for 5 min at RT while rocking. The Pierce ECL Western Blotting Substrate (Fisher #32209) was used for detection according to manufacturer instructions. The nitrocellulose membrane was imaged with the ImageQuant™ LAS 4000 (GE Healthcare). Quantification was done using the open image analysis software ImageJ. Relative abundance of phospho-STAT1 was normalized to the loading control protein alpha-Tubulin.

### RNA isolation, cDNA synthesis, and q-RT-PCR

Cells were harvested at 0, 6, 12, and 24 h post IFN treatment, and RNA was isolated using RNAeasy RNA extraction kit (Qiagen) as per manufacturer’s instructions. DNA was synthesized using iSCRIPT reverse transcriptase (BioRad) from 250 ng of total RNA per 20 µL reaction according to the manufacturer’s instructions. Quantitative RT-PCR assay was performed using iTaq SYBR green (BioRad) as per manufacturer’s instructions. The expression of the various ISGs was normalized to the housekeeping gene *TBP.* The expression levels of the various ISG were then normalized to mock of each time-point, to obtain the fold change expression to mock treated cells. Primer sequences are listed below.

**Table 1.**
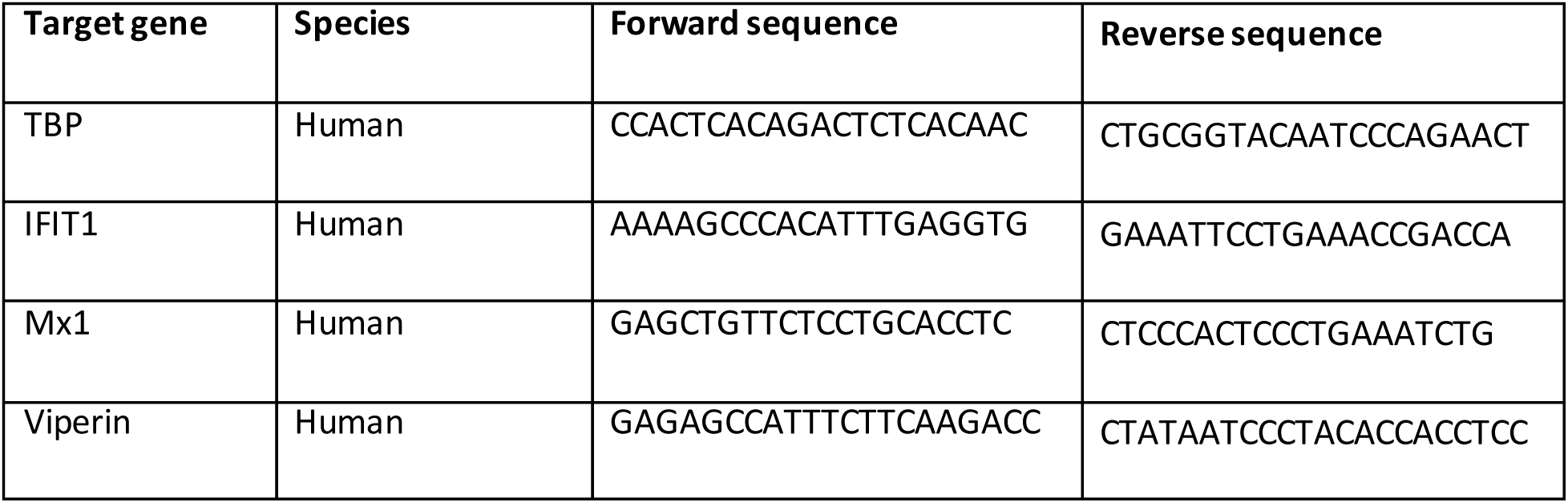
Primer sequences for RT-q-PCR.

### Analysis of spatial heterogeneity using image analysis software

T84 prom-Mx1-eGFP cells seeded on glass bottom 8-well chamber slides (iBIDI) were mock treated or treated with IFNs for 24 h and fixed in 2% paraformaldehyde (PFA) (in PBS) for 20 mins at RT. Cells were washed in 1X PBS and permeabilized in 0.5% Triton-X-100 (in PBS) for 15 mins at RT. Cell nuclei were stained with DAPI (BD Biosciences #564907) diluted 1:1000 in PBS for 20 mins. Cells were washed in 1X PBS three times and maintained in PBS. Cells were imaged on a ZEISS Celldiscoverer 7 Widefield microscope using a 20 x 0.5 magnification (Numerical Aperture NA = 0.5).

To analyze the spatial heterogeneity of IFN-dependent immune response, we first generated masks from DAPI images representing each nucleus as an individual object with the segmentation software Ilastik 1.2.0. These masks were then used in CellProfiler 3.1.9 to determine (a) the XY-localization of each object (nucleus) within its 2-dimensional plane and (b) to measure the prom-Mx1-eGFP fluorescence intensity within each object (nucleus). Using the information on the XY-localization, we applied the DBSCAN-CellX-App (https://github.com/GrawLab/DBSCAN-CellX/) to the data to assess whether a cell is localized at the edge or the center of a cluster, and to determine the edge degree of a cell. The cell localization and cell edge degree were plotted against the percentage of prom-Mx1-eGFP positive cells (as compared to the mock-treated samples) within each sub-population group, resulting in the visualization of the immune response of single cells within their population context.

### Surface micropatterning, cell seeding and image analysis

For glass micropatterning by Quartz mask-based approach (ultraviolet light-Ozone (UVO)-based micropatterning of glass surfaces using a Quartz-mask), a quartz chromium photomask containing 200 µm diameter clear circles was custom made by Toppan Photomasks Inc. (Mask type = 1X Master, mask size = 4” x 4” x 0.06”). The UVO-based micropatterning protocol was adapted from Pitaval et al. (51). Briefly, glass coverslips of 25 mm diameter (Marienfeld # 0117650) were pre-cleaned with 100% ethanol for 15 min while sonicating, rinsed twice with deionized water, and dried with compressed air. The glass coverslips were activated in the UVO-Cleaner® Model 30 (Jelight Company Inc.) for 10 min and then passivated for 45 min at room temperature with 100 µl 0.1 mg/ml poly-L lysin/poly-ethylene glycol PLL(20)-g[3.5]-PEG(2) (SuSoS Surface Technology) in water. After passivation, the coverslips were washed twice with deionized water for 10 min. Before the micropatterning step, the photomask was washed with acetone and isopropanol, dried with a stream of compressed air and cleaned in the UVO cleaner for 5 min. Directly after cleaning, the passivated glass coverslips were sandwiched with the photomask using 8 µl of deionized water to create an intimate contact between the chromium side of the photomask and the passivated surface of the coverslip. The photomask with the coverslips was placed in the UVO cleaner (quartz side facing towards UVO light) for 5 min for the micropatterning step. After UVO exposure, the coverslips were carefully detached from the photomask and stored in PBS at 4°C until further use.

Transwell micropatterning using maskless photolithography system (Supp. Fig. 4): 6.5 mm Transwell® with 3.0 µm Pore Polycarbonate Membrane Inserts (Costar #CLS3415) were used. The transwell membrane was activated in the plasma cleaner (Tepla 100-E Plasma System) at 0.4 mbar O_2_-pressure and 200 W for 1 min. The surface was then incubated with 0.1% (w/v) Poly-L-Lysin (PLL) solution (in H_2_O) (Sigma #P8920) for 30 mins at RT, washed four times with deionized water and dried with compressed air. The surface was passivated with 90 µl 90 mg/ml Methoxy-Poly (Ethylene Glycol)-Succinimidyl Valerate (mPEG-SVA) (5000Da) (Laysan Bio Inc. #MPEG-SVA-5000) in 0.1 M HEPES buffer (pH 8.4) for 1 h at RT. During this reaction the SVA ester covalently binds to the amines of the PLL, resulting in a homogenous passivation of the glass surface with a PLL-PEG polymer. The surface was washed four times with deionized water and dried with compressed air. 0.5 µl photoactivator PLPP-gel (Alvéole Lab, www.alveolelab.com) was put in the center of the surface. Immediately after, 16 µl of 100% EtOH were added on the top of the PLPP-gel and the mixture was homogenized by manual rotation and the surface was dried at RT. This system is able to micropattern any previously designed pattern on any surface. To design a pattern, the open-source software Inkscape (inkscape.org) was used with the following scale: 1 px corresponded to 0.28 µm. We designed circles with 200 µm diameter, which was then loaded into the Leonardo software (Alvéole Lab) for micropatterning. The micropatterning was performed on a Nikon Eclipse Ti2 Microscope with a 20x S Plan Fluor ELWD Objective (NA = 0.45). The passivated surface coated with the photoactivator PLPP was placed on the microscope stage. The photo-micropatterning was controlled with the Leonardo software and executed by the PRIMO optical module (Alvéole Lab) using the stitching mode and a 375 nm laser at a dose of 30 mJ/mm^2^. The patterned surface was then washed six times with deionized water and stored in PBS at 4°C until further use.

For cell seeding, the patterned surface was coated with 2.5% human collagen (Sigma #C5533-5MG) diluted in water for 1 h at RT. An excess of T84 pMx1-mCherry or T84 pMx1-eGFP cells were then seeded and incubated for 2 h at 37°C. Precisely, 1,000,000 cells in 2 ml culturing media were used for the 25 mm diameter coverslips (in a 6-well cell culture plate Greiner Bio-One #657160) and 200,000 cells in 200 µl culturing media were added to the apical compartment of transwell (the basolateral compartment of transwells was filled with 600 µl culturing media). Non-adherent cells were then washed away 2 h post seeding with 2 times PBS, and then fresh culturing medium was added. One day post-seeding, a medium change was performed, and on the second day post-seeding cells were treated with IFNs.

For cells seeded on micropatterned 25 mm diameter coverslips: Immediately after treating, live cell imaging was performed using a ZEISS Celldiscoverer 7 Widefield microscope using a 20 x 1 magnification (NA = 0.8) at 37°C and 5% CO_2_ for 24 h, taking an image every 12 h starting at 0 h post treatment. For cells seeded on micropatterned transwell: Cells were fixed in 2% PFA at 0, 12, and 24 h post treatment and mounted with DAPI (Invitrogen, #P36935). Imaging was performed using the ZEISS Celldiscoverer 7 Widefield microscope using a 20 x 1 magnification (NA = 0.8). To analyze the spatial heterogeneity of IFN-dependent immune response, CellProfiler 3.1.9 was used to generate masks that divide each population into an edge and center, and to measure the prom-Mx1-eGFP fluorescence intensity within the edge or the center of the population. The fluorescence intensity was normalized to the 0 h time-point as described in the figure legends.

### Surface biotinylation and surface proteome analysis

T84 cells were cultured in Corning transwell inserts (1.5*10^5^ cells/insert) in 1% O_2_ atmosphere until fully polarized (typically 2-4 days). Cells were then washed thrice in PBS and treated with 1 mg/ml Sulfo NHS-SS-Biotin (ThermoFisher) in biotinylation buffer (10 mM HEPES, 130 mM NaCl, 2 mM MgSO_4_, 1 mM CaCl_2_, pH 7.9) on the apical or basolateral side for 15 minutes on ice (biotinylation buffer was added to the opposite side to prevent drying). After incubation, cells were washed with 100 mM glycine for three times (last wash was left on cells for 10 minutes) to remove and quench excess biotin. Membranes were then cut and added to cell lysis buffer (50 mM HEPES, 150 mM NaCl, 5 mM EDTA, 1% Triton x-100, 0.1% SDS, pH 7.4) for 30 minutes in ice. After brief sonication and sedimentation of insoluble fragments, the protein amount was quantified using the DC protein assay kit (Biorad) and the same amount of total lysate for each sample (apical, basolateral and no biotinylation sample (background)) was loaded with High Capacity Neutravidin Agarose beads (ThermoFisher) and incubated overnight at 4°C on an orbital shaker. The day after, supernatant was removed and beads were washed twice with high salt buffer (1M NaCl, 50 mM HEPES, 0.1% Triton x-100, pH 7.4) and twice in 50 mM HEPES (pH 7.4). Beads were then incubated in Laemmli buffer for 20 minutes at RT on an orbital shaker. The supernatant containing the biotinylated surface proteins was then harvested, and loaded and ran on an SDS-PAGE for purification. Bands were excised and digested wit trypsin using a standard protocol (52). After digestion, peptides were extracted and dried for LC-MS analysis. Peptides were reconstituted in 15 µl of 0.05% trifluoroacetic acid, 4% acetonitrile, and 6.6 µl were analyzed by an Ultimate 3000 reversed-phase capillary nano liquid chromatography system connected to a Q Exactive HF mass spectrometer (Thermo Fisher Scientific). Samples were injected and concentrated on a trap column (PepMap100 C18, 3 µm, 100 Å, 75 µm i.d. x 2 cm, Thermo Fisher Scientific) equilibrated with 0.05% trifluoroacetic acid in water. LC separations were performed on a capillary column (Acclaim PepMap100 C18, 2 µm, 100 Å, 75 µm i.d. x 25 cm, Thermo Fisher Scientific) at an eluent flow rate of 300 nl/min. Mobile phase A contained 0.1 % formic acid in water, and mobile phase B contained 0.1% formic acid in 80 % acetonitrile / 20% water. The column was pre-equilibrated with 5% mobile phase B followed by an increase of 5-44% mobile phase B in 100 min. Mass spectra were acquired in a data-dependent mode utilising a single MS survey scan (m/z 350–1650) with a resolution of 60,000 and MS/MS scans of the 15 most intense precursor ions with a resolution of 15,000. The dynamic exclusion time was set to 20 seconds and automatic gain control was set to 3×10^6^ and 1×10^5^ for MS and MS/MS scans, respectively.

MS and MS/MS raw data were analyzed using the MaxQuant software package (version 1.6.14.0) with implemented Andromeda peptide search engine (53). Data were searched against the human reference proteome downloaded from Uniprot (75,074 sequences, taxonomy 9606, last modified March 10, 2020) using the default parameters except for the following changes: label-free quantification (LFQ) enabled, match between runs enabled, iBAQ enabled, max missed cleavages: 3.

Perseus downstream analysis was performed as follows: Proteins were cross referenced with the UniProt human database for gene ontology terms (Plasma membrane, plasma membrane part, cell surface, cell outer membrane), then filtered out if they had less than 3 replicates or if they had no GO term matching the above mentioned search. Background samples were used to filter out any protein nonspecifically bound to the Neutravidin beads. Apical/Basolateral ratios were calculated based on the apical LFQ signal divided by the total LFQ signal. Pairwise t tests were conducted to determine significant differential protein expression.

### FITC-Dextran permeability assay

T84 cells were grown on transwell inserts as a monolayer, with 600 µL media in the basolateral compartment and 200 µL media in the apical compartment. Media was removed from the apical compartment of the transwell and replaced by 200µL of fresh medium containing of 2 mg/mL fluorescein isothiocyanate (FITC)-labelled dextran (4 kDa) (Sigma-Aldrich, # 46944-500MG-F). As a negative control and to calculate the background, culture media alone was used on a well without cells. For the positive control (maximum diffusion of FITC-Dextran from apical to basolateral compartment) 200 µL of 2 mg/mL of FITC-Dextran was added to the apical side of a well without cells and 600µL culturing media were added to the basolateral compartment. Cells and controls were incubated for 3 h at 37°Cand then media was collected from the basolateral compartment. Fluorescent signal was measured using an 800TS Microplate Reader (BioTek) at an excitation wavelength of 495nm. A standard curve by serial dilution of the FITC-Dextran in culturing media was done to assess the basolateral FITC-Dextran concentration.

### Viral infections

All virus infections were performed with a multiplicity of infection (MOI) of 1 as determined in T84 cells. Cells were mock treated or pre-treated with type I and type III IFNs for 24 h. Pre-treatment culture medium was replaced with fresh medium containing MRV or VV at an MOI of 1 and supplemented with the same interferons as the pre-treatment. 16 h post-infection, cells were fixed in 2% PFA for immunofluorescent staining.

### Indirect immunofluorescence assay

Cells were seeded on glass coverslips for ZO1 staining and on plastic bottom multiwells for viral protein staining. Cells were fixed in 2% paraformaldehyde (PFA) for 20 mins at RT. Cells were washed in 1X PBS and permeabilized in 0.5% Triton-X100 diluted in PBS for 15 mins at RT. Cells were blocked using 3% BSA in PBS for 30 min at RT. Antibodies against ZO1 (Thermo Fisher Scientific #40-2200) or Non-Structural Mammalian Reovirus Protein µNS (54) were diluted in 1% BSA (in PBS) and incubated for 1 h at RT. Cells were washed with PBS three times, and incubated with DAPI (BD Biosciences, #564907) and secondary antibody conjugated to AF488 (Molecular Probes) diluted 1:1000 in 1% BSA (in PBS) for 30 mins at RT. For cells infected with Vaccinia virus, immunostaining was omitted since the virus strains expresses eGFP and only the DAPI staining was done for visualization of cell nuclei. Cells were washed in 1X PBS three times. Cells seeded on coverslips were mounted, and cells seeded in multiwells were maintained in PBS until imaging. Cells were imaged on a ZEISS Celldiscoverer 7 Widefield microscope using a 20 x 0.5 magnification (Numerical Aperture NA = 0.5).

### Statistics and computational analyses

All statistical analyses were performed by statistical tests as specified in figure legends using the GraphPad Prism software package (Version 8.0.1).

To quantify the number of Vaccinia virus eGFP or MRV infected cells, ilastik 1.2. 0 was used on DAPI images to generate a mask representing each nucleus as an individual object. These masks were used on CellProfiler 3.1.9 to measure the fluorescence intensity coming from the virus infection within each nucleus. A threshold was set based on the basal fluorescence of non-infected samples, and all nuclei with a higher fluorescence were counted as infected cells.

## Acknowledgements

SB and MS are supported by the UF College of Medicine start-up package funds. This project was supported by the Deutsche Forschungsgemeinschaft (DFG) by the projects SFB 1129 (DFG 240245660) and TRR186 (DFG 278001972). TH was supported by SFB 1129 TP11 (DFG 240245660). FG was supported by the Chica and Heinz Schaller Foundation. PD was supported by Deutscher Akademischer Austauschdienst (DAAD). For mass spectrometry, we would like to acknowledge the assistance of the Core Facility BioSupraMol supported by the DFG. We thank Joel Christian for introducing us to the maskless photolithography micropatterning system (Alvéole Lab, www.alveolelab.com). We thank the Max Planck Society for its support and thank the Boulant and Stanifer lab members for the constructive discussions and for proofreading this manuscript.

## Expanded View Figure Legends

**Supplementary Figure 1: IFN-sensing reporter cell lines shows single cell behavior during IFN treatment.** (A) Schematic depicting the T84 prom-Mx1-fp reporter cell line. Upon interaction of IFNs with their receptor, downstream signaling induces nuclear translocation of the transcription complex ISGF3. This leads to expression of the fluorescent protein under control of the ISG Mx1 promoter. The fluorescent protein accumulates in the cytosol and can be visualized by fluorescence microscopy. (B) Representative images showing expression of the fluorescent reporter (white) after mock, 2000 IU/mL IFNβ1, or 300 ng/mL IFNλ1-3 treatment. Nuclei are stained with DAPI (blue). n=3 biological replicates. Scale bar = 100µm.

**Supplementary Figure 2: Temporal response of cells at high and low density to IFN treatment.** T84-prom-Mx1-fp cells at high or low density were treated with increasing concentrations of (A) IFNβ1 and (B) IFNλ1-3. Live cell fluorescence imaging was performed at an interval of 2 h for 24 h. The mean fluorescence intensity (MFI) of the reporter expression within each cell was averaged for each density and normalized to the mock MFI of each time-point (fold change). n = 3 biological replicates. n.s =not significant. P<0.05 *, P<0.01 **, P<0.001 ***, P <0.0001 **** as determined by Unpaired t test with Welch’s correction.

**Supplementary Figure 3: Transwell system allows for the formation of a semipermeable monolayer of polarized cells.** T84 cells were seeded on transwell inserts to allow for a polarized monolayer formation. (A) 5 days post seeding, cells were fixed, and indirect immunofluorescence was performed against the junctional complex protein ZO1 (green). Nuclei were stained with DAPI (blue). Representative image is shown. Scale bars=50 µm. n=3 biological replicates. (B) Formation and integrity of the monolayer was followed by measuring the transepithelial electrical resistance (TEER) (Ω/cm^2^) over 5 days. Values > 1000 Ω/cm^2^ (dotted line) shows that cells established a polarized monolayer formation. n = 3 biological replicates. (C) 5 days post seeding, after reaching a polarized monolayer, the integrity of the monolayer was confirmed by the FITC-Dextran permeability assay. Diffusion of FITC-Dextran from the apical to the basolateral compartment was measured and expressed as concentration (mg/mL) of FITC-Dextran in the basolateral compartment after 3h incubation. Positive control (pos) was the maximum diffusion possible and the negative control (neg) was medium only without FITC-Dextran. n ≥ 3 biological replicates. n.s. =not significant, P<0.05 *, P<0.01 **, P<0.001 ***, P <0.0001 **** as determined by ordinary one-way ANOVA with Dunnett’s multiple comparison test using the positive control as reference.

**Supplementary Figure 4: Micropatterning of transwell inserts.** Schematic depicting the micropatterning on transwell membranes using the PRIMO system (Alvéole Lab, www.alveolelab.com).

## References

1. Kawai T, Akira S. 2006. Innate immune recognition of viral infection. Nat Immunol. Nature Publishing Group.

2. Koyama S, Ishii KJ, Coban C, Akira S. 2008. Innate immune response to viral infection. Cytokine. Cytokine.

3. Novick D, Cohen B, Rubinstein M. 1994. The human interferon α β receptor: Characterization and molecular cloning. Cell 77:391–400.

4. Kotenko S V., Gallagher G, Baurin V V., Lewis-Antes A, Shen M, Shah NK, Langer JA, Sheikh F, Dickensheets H, Donnelly RP. 2003. IFN-λs mediate antiviral protection through a distinct class II cytokine receptor complex. Nat Immunol. Nat Immunol.

5. Sheppard P, Kindsvogel W, Xu W, Henderson K, Schlutsmeyer S, Whitmore TE, Kuestner R, Garrigues U, Birks C, Roraback J, Ostrander C, Dong D, Shin J, Presnell S, Fox B, Haldeman B, Cooper E, Taft D, Gilbert T, Grant FJ, Tackett M, Krivan W, McKnight G, Clegg C, Foster D, Klucher KM. 2003. IL-28, IL-29 and their class II cytokine receptor IL-28R. Nat Immunol. Nat Immunol.

6. Sommereyns C, Paul S, Staeheli P, Michiels T. 2008. IFN-Lambda (IFN-λ) Is Expressed in a Tissue-Dependent Fashion and Primarily Acts on Epithelial Cells In Vivo. PLoS Pathog 4:e1000017.

7. Pott J, Mahlakõiv T, Mordstein M, Duerr CU, Michiels T, Stockinger S, Staeheli P, Hornef MW. 2011. IFN-λ determines the intestinal epithelial antiviral host defense. Proc Natl Acad Sci U S A 108:7944–7949.

8. Mordstein M, Neugebauer E, Ditt V, Jessen B, Rieger T, Falcone V, Sorgeloos F, Ehl S, Mayer D, Kochs G, Schwemmle M, Günther S, Drosten C, Michiels T, Staeheli P. 2010. Lambda Interferon Renders Epithelial Cells of the Respiratory and Gastrointestinal Tracts Resistant to Viral Infections. J Virol 84:5670–5677.

9. Levy DE, Marié IJ, Durbin JE. 2011. Induction and function of type i and III interferon in response to viral infection. Curr Opin Virol. Elsevier B.V.

10. Schindler C, Levy DE, Decker T. 2007. JAK-STAT signaling: From interferons to cytokines. J Biol Chem. J Biol Chem.

11. Stanifer ML, Pervolaraki K, Boulant S. 2019. Differential regulation of type I and type III interferon signaling. Int J Mol Sci. MDPI AG.

12. Altschuler SJ, Wu LF. 2010. Cellular Heterogeneity: Do Differences Make a Difference? Cell. Elsevier B.V.

13. Spencer SL, Gaudet S, Albeck JG, Burke JM, Sorger PK. 2009. Non-genetic origins of cell-to-cell variability in TRAIL-induced apoptosis. Nature 459:428–432.

14. Tay S, Hughey JJ, Lee TK, Lipniacki T, Quake SR, Covert MW. 2010. Single-cell NF-B dynamics reveal digital activation and analogue information processing. Nature 466:267–271.

15. Roesch A, Fukunaga-Kalabis M, Schmidt EC, Zabierowski SE, Brafford PA, Vultur A, Basu D, Gimotty P, Vogt T, Herlyn M. 2010. A Temporarily Distinct Subpopulation of Slow-Cycling Melanoma Cells Is Required for Continuous Tumor Growth. Cell 141:583–594.

16. Gupta PB, Fillmore CM, Jiang G, Shapira SD, Tao K, Kuperwasser C, Lander ES. 2011. Stochastic state transitions give rise to phenotypic equilibrium in populations of cancer cells. Cell 146:633–644.

17. Slack MD, Martinez ED, Wu LF, Altschuler SJ. 2008. Characterizing heterogeneous cellular responses to perturbations. Proc Natl Acad Sci U S A 105:19306–19311.

18. Sharma S V., Lee DY, Li B, Quinlan MP, Takahashi F, Maheswaran S, McDermott U, Azizian N, Zou L, Fischbach MA, Wong KK, Brandstetter K, Wittner B, Ramaswamy S, Classon M, Settleman J. 2010. A Chromatin-Mediated Reversible Drug-Tolerant State in Cancer Cell Subpopulations. Cell 141:69–80.

19. Patil S, Fribourg M, Ge Y, Batish M, Tyagi S, Hayot F, Sealfon SC. 2015. Single-cell analysis shows that paracrine signaling by first responder cells shapes the interferon-β response to viral infection. Sci Signal 8.

20. Rand U, Rinas M, Werk JS, Nöhren G, Linnes M, Kröger A, Flossdort M, Kály-Kullai K, Hauser H, Höfer T, Köster M. 2012. Multi-layered stochasticity and paracrine signal propagation shape the type-l interferon response. Mol Syst Biol 8.

21. Zhao M, Zhang J, Phatnani H, Scheu S, Maniatis T. 2012. Stochastic expression of the interferon-β gene. PLoS Biol 10.

22. Wimmers F, Subedi N, van Buuringen N, Heister D, Vivié J, Beeren-Reinieren I, Woestenenk R, Dolstra H, Piruska A, Jacobs JFM, van Oudenaarden A, Figdor CG, Huck WTS, de Vries IJM, Tel J. 2018. Single-cell analysis reveals that stochasticity and paracrine signaling control interferon-alpha production by plasmacytoid dendritic cells. Nat Commun 9.

23. Schmid B, Rinas M, Ruggieri A, Acosta EG, Bartenschlager M, Reuter A, Fischl W, Harder N, Bergeest J-P, Flossdorf M, Rohr K, Höfer T, Bartenschlager R. 2015. Live Cell Analysis and Mathematical Modeling Identify Determinants of Attenuation of Dengue Virus 2’-O-Methylation Mutant. PLOS Pathog 11:e1005345.

24. Maier BD, Aguilera LU, Sahle S, Mutz P, Kalra P, Dächert C, Bartenschlager R, Binder M, Kummer U. 2022. Stochastic dynamics of Type-I interferon responses. PLoS Comput Biol 18.

25. Shalek AK, Satija R, Adiconis X, Gertner RS, Gaublomme JT, Raychowdhury R, Schwartz S, Yosef N, Malboeuf C, Lu D, Trombetta JJ, Gennert D, Gnirke A, Goren A, Hacohen N, Levin JZ, Park H, Regev A. 2013. Single-cell transcriptomics reveals bimodality in expression and splicing in immune cells. Nature 498:236–240.

26. Andrews SS, Dinh T, Arkin AP. 2009. Stochastic Models of Biological Processes, p. 8730–8749. In Encyclopedia of Complexity and Systems Science. Springer New York.

27. Maier BD, Aguilera LU, Sahle S, Mutz P, Kalra P, Dächert C, Bartenschlager R, Binder M, Kummer U. 2022. Stochastic dynamics of Type-I interferon responses. PLoS Comput Biol 18.

28. Snijder B, Pelkmans L. 2011. Origins of regulated cell-to-cell variability. Nat Rev Mol Cell Biol 12:119–125.

29. Zeng L, Skinner SO, Zong C, Sippy J, Feiss M, Golding I. 2010. Decision Making at a Subcellular Level Determines the Outcome of Bacteriophage Infection. Cell 141:682–691.

30. St-Pierre F, Endy D. 2008. Determination of cell fate selection during phage lambda infection. Proc Natl Acad Sci U S A 105:20705–20710.

31. Robert L, Paul G, Chen Y, Taddei F, Baigl D, Lindner AB. 2010. Pre-dispositions and epigenetic inheritance in the Escherichia coli lactose operon bistable switch. Mol Syst Biol 6.

32. Snijder B, Sacher R, Rämö P, Damm E-M, Liberali P, Pelkmans L. 2009. Population context determines cell-to-cell variability in endocytosis and virus infection. Zurich PhD Progr Mol Life Sci 461.

33. Kavaliauskiene S, Nymark CM, Bergan J, Simm R, Sylvänne T, Simolin H, Ekroos K, Skotland T, Sandvig K. 2014. Cell density-induced changes in lipid composition and intracellular trafficking. Cell Mol Life Sci 71:1097–1116.

34. Trajkovic K, Valdez C, Ysselstein D, Krainc D. 2019. Fluctuations in cell density alter protein markers of multiple cellular compartments, confounding experimental outcomes. PLoS One 14.

35. Chelakkot C, Ghim J, Ryu SH. 2018. Mechanisms regulating intestinal barrier integrity and its pathological implications. Exp Mol Med. Nature Publishing Group.

36. Bhushal S, Wolfsmüller M, Selvakumar TA, Kemper L, Wirth D, Hornef MW, Hauser H, Köster M. 2017. Cell polarization and epigenetic status shape the heterogeneous response to type III interferons in intestinal epithelial cells. Front Immunol 8.

37. Pervolaraki K, Rastgou Talemi S, Albrecht D, Bormann F, Bamford C, Mendoza JL, Garcia KC, McLauchlan J, Höfer T, Stanifer ML, Boulant S. 2018. Differential induction of interferon stimulated genes between type I and type III interferons is independent of interferon receptor abundance. PLoS Pathog 14.

38. Zambarda C, Pérez González C, Schoenit A, Veits N, Schimmer C, Jung R, Ollech D, Christian J, Roca-Cusachs P, Trepat X, Cavalcanti-Adam EA. 2022. Epithelial cell cluster size affects force distribution in response to EGF-induced collective contractility. Eur J Cell Biol 101:151274.

39. Doldan P, Dai J, Metz-Zumaran C, Patton JT, Stanifer ML, Boulant S. 2022. Type III and Not Type I Interferons Efficiently Prevent the Spread of Rotavirus in Human Intestinal Epithelial Cells. J Virol 96.

40. Cao X, Surma MA, Simons K. 2012. Polarized sorting and trafficking in epithelial cells. Cell Res. Cell Res.

41. Hartsock A, Nelson WJ. 2008. Adherens and tight junctions: Structure, function and connections to the actin cytoskeleton. Biochim Biophys Acta - Biomembr. Elsevier.

42. Benson K, Cramer S, Galla HJ. 2013. Impedance-based cell monitoring: Barrier properties and beyond. Fluids Barriers CNS. Fluids Barriers CNS.

43. Schmid B, Rinas M, Ruggieri A, Acosta EG, Bartenschlager M, Reuter A, Fischl W, Harder N, Bergeest JP, Flossdorf M, Rohr K, Höfer T, Bartenschlager R. 2015. Live Cell Analysis and Mathematical Modeling Identify Determinants of Attenuation of Dengue Virus 2’-O-Methylation Mutant. PLoS Pathog 11.

44. Fang Y, Sullivan R, Graham CH. 2007. Confluence-dependent resistance to doxorubicin in human MDA-MB-231 breast carcinoma cells requires hypoxia-inducible factor-1 activity. Exp Cell Res 313:867–877.

45. Meli L, Jordan ET, Clark DS, Linhardt RJ, Dordick JS. 2012. Influence of a three-dimensional, microarray environment on human Cell culture in drug screening systems. Biomaterials 33:9087–9096.

46. Jaspers I, Ciencewicki JM, Brighton LE. 2009. Localization of type i interferon receptor limits interferon-induced TLR3 in epithelial cells. J Interf Cytokine Res 29:289–297.

47. Humlicek AL, Manzel LJ, Chin CL, Shi L, Excoffon KJDA, Winter MC, Shasby DM, Look DC. 2007. Paracellular Permeability Restricts Airway Epithelial Responses to Selectively Allow Activation by Mediators at the Basolateral Surface. J Immunol 178:6395–6403.

48. Stanifer ML, Rippert A, Kazakov A, Willemsen J, Bucher D, Bender S, Bartenschlager R, Binder M, Boulant S. 2016. Reovirus intermediate subviral particles constitute a strategy to infect intestinal epithelial cells by exploiting TGF-β dependent pro-survival signaling. Cell Microbiol 18:1831–1845.

49. Mercer J, Helenius A. 2008. Vaccinia virus uses macropinocytosis and apoptotic mimicry to enter host cells. Science (80-) 320:531–535.

50. Cotter CA, Earl PL, Wyatt LS, Moss B. 2015. Preparation of Cell Cultures and Vaccinia Virus Stocks. Curr Protoc Microbiol 39:14A.3.1–14A.3.18.

51. Pitaval A, Tseng Q, Bornens M, Théry M. 2010. Cell shape and contractility regulate ciliogenesis in cell cycle-arrested cells. J Cell Biol 191:303–312.

52. Shevchenko A, Wilm M, Vorm O, Mann M. 1996. Mass spectrometric sequencing of proteins silver-stained polyacrylamide gels. Anal Chem 68:850–858.

53. Tyanova S, Temu T, Cox J. 2016. The MaxQuant computational platform for mass spectrometry-based shotgun proteomics. Nat Protoc 11:2301–2319.

54. Shah PNM, Stanifer ML, Höhn K, Engel U, Haselmann U, Bartenschlager R, Kräusslich HG, Krijnse-Locker J, Boulant S. 2017. Genome packaging of reovirus is mediated by the scaffolding property of the microtubule network. Cell Microbiol 19.

